# ZIP13 regulates lipid metabolism by changing intracellular iron and zinc balance

**DOI:** 10.1101/2024.07.03.601606

**Authors:** Ayako Fukunaka, Gen Tanaka, Toru Kimura, Kowit Hengphasatporn, Daisuke Saito, Mayo Shigeta, Hiroshi Shibata, Takayuki Ichinose, Ichitaro Abe, Mari Shimura, Takashi Sato, Yuko Nakagawa, Takuro Horii, Izuho Hatada, Shingo Kajimura, Hiroshi Kiyonari, Hiroyuki Sakurai, Yasuteru Shigeta, Hirotaka Watada, Toshiyuki Fukada, Yoshio Fujitani

**Author notes:** **Corresponding author** (T.F.), (A.F.), and (Y.F.). These authors contributed equally to this work (Ayako Fukunaka, Gen Tanaka, Toru Kimura).

## Abstract

Metabolic diseases are caused by a prolonged energy imbalance, and adipose tissue is known to be the main contributor. We previously reported that ZIP13, an Slc39a transporter whose deficiency causes Ehlers-Danlos syndrome spondylocheirodysplastic type 3 associated with lipoatrophy, inhibits the adipocyte browning pathway by modulating intracellular zinc status. The precise mechanisms of how ZIP13 regulates the homeostasis of adipose tissue remain unclear and therefore, we investigated the role of ZIP13 in mature adipocytes using adipocyte-specific *Zip13*-deficient mice. We herein demonstrate that these mice show accelerated lipolysis and reduced respiratory exchange ratio. In addition, abundance of iron and zinc balance were altered during differentiation in normal adipocytes, whereas iron distribution was substantially affected in *Zip13*-deficient adipocytes, which downregulated PDE activity and enhanced β-adrenergic receptor signaling pathways. Importantly, we confirmed that ZIP13 could transport both zinc and iron, using the *Xenopus* oocyte transport system and *in silico* structural dynamics simulations, and that the defect in iron distribution perturbs proper lipolysis. Together, these results illustrate that ZIP13 acts as a key regulator for lipolysis in adipocytes via the proper use of metals, and that the ZIP13-iron axis plays an important role in regulation of lipid metabolism.

## Introduction

Obesity and its associated metabolic diseases are caused by a long-term imbalance between energy intake and energy expenditure. Adipose tissue, which is a major factor in controlling the energy balance, is composed of white and brown adipocytes, which have two distinct functions; i.e., white adipocytes store excess energy, whereas brown adipocytes specialize in consuming energy ^1, 2^. As chronic caloric excess is a main drivers of the development of obesity and type 2 diabetes, most studies have mainly focused on macronutrients, such as the effects of the types or compositions of fats and carbohydrates on metabolic diseases. However, little is known about the role of micronutrients and the proteins that regulate the homeostasis of each micronutrient in obesity and type 2 diabetes ^3, 4, 5, 6, 7, 8^. Therefore, in this study, we aimed to investigate how essential trace elements are involved in metabolic homeostasis and events, by focusing on proteins associated with iron, zinc, and micronutrients, such metal ion transporters.

Zrt/Irt proteins (ZIPs and SLC39A) consist of a family of divalent metal ion transporters. Zrt stands for zinc-regulated transporter, which was named after identification of the yeast zinc importer, and Irt stands for iron-regulated transporter after the plant iron importer was discovered ^9^. Although some ZIP members selectively transport zinc, as can be inferred from the name of ZIP, other members, such as ZIP8 and ZIP14, have broader substrate specificity and transport divalent metal ions, including zinc, iron, manganese, and cadmium *in vitro* ^10, 11^. Genetic approaches involving gene-knockout (KO) mice and human diseases indicate that the physiologically transported substrate of ZIP8 and ZIP14 *in vivo* is manganese ions ^12, 13, 14^. ZIP13 is an intracellular ZIP member that has been reported to be localized to the Golgi, which was demonstrated to predominantly transport zinc ions by a competition assay using other metals, including iron ^15^. We and other groups have also demonstrated that zinc levels are altered upon either ectopic expression or downregulation of ZIP13, all of which were analyzed using zinc-specific probes or by measuring metallothionein (*MT*) expression ^15, 16, 17, 18^, because ZIP13 localizes to the intracellular compartment which makes it difficult to elucidate whether ZIP13 can transport the zinc, or other metals, *in vivo* and *in vitro*.

Among the ZIP family members, ZIP8, ZIP14 and ZIP13 have been reported to participate in metabolic regulation, and hence their dysfunctions are associated with diseases, such as obesity, abnormal high density lipoprotein cholesterol levels, lipodystrophy, etc. ^19, 20, 21^. We and other group previously demonstrated that ZIP13 plays an important role in the development of connective tissue in mice and humans, and its dysfunction causes Ehlers-Danlos syndrome spondylocheirodysplastic type 3(EDSSPD3, OMIM 612350), which is a very rare autosomal recessive disease in which patients develop lipoatrophy, strongly indicating the crucial role of ZIP13 in fat tissues homeostasis ^21, 22^. We also reported that *Zip13* KO preferentially promotes beige adipocyte biogenesis and energy expenditure, thereby ameliorating diet-induced obesity and insulin resistance ^18^, although the precise molecular mechanisms by which ZIP13 controls adipocytes functions remain unclear.

Here, we investigated the role of ZIP13 in mature adipocytes, using adipocyte-specific *Zip13* KO (Adipo-*Zip13*KO) mice showed accelerated lipolysis in adipocytes leading to reduced respiratory exchange ratios (RER). We also found that abundance of iron and zinc balance was altered during differentiation in control adipocytes, whereas iron distribution was substantially affected in *Zip13*-deficient adipocytes, which downregulated PDE activity and enhanced β-adrenergic receptor signaling pathways. Furthermore, we confirmed that ZIP13 can transport both zinc and iron, using the *Xenopus* oocyte transport system and *in silico* structural dynamics simulations, and that ZIP13-mediated intracellular iron homeostasis is crucial for proper lipid metabolism. Together, these results indicate that ZIP13 regulates lipolysis in adipocytes via the proper use of metals, demonstrating the ZIP13-iron axis plays an important role in regulation of lipid metabolism.

## Results

### Adipo-*Zip13*KO mice demonstrate accelerated lipolysis in a cell-autonomous manner

To investigate the intrinsic role of ZIP13 in mature adipocytes, we generated adipocyte-specific *Zip13* KO mice by crossing *Zip13flox/flox;* adiponectin-Cre mice with *Zip13*+/-mice (hereafter referred to as Adipo-*Zip13*KO mice), in which we confirmed that *Zip13* mRNA expression was reduced in all fat tissues analyzed (Fig. 1A). Adipo-*Zip13*KO mice appeared to be resistant to an increase in body weight under HFD conditions (Fig. 1B and C), similar to systemic *Zip13* KO mice^18^. We also found that the body weights of Adipo-*Zip13*KO mice were decreased even in the standard diet (STD) condition in mice older than 30 weeks (Fig. 1D), together with a reduction in adipose tissues weight, particularly in subcutaneous fat tissue (Fig. 1E, and Figs. S1A and B). Intriguingly, Adipo-*Zip13*KO mice appeared to have normal glucose tolerance, but their insulin concentrations were lower than those of Ctrl mice under STD condition (Fig. S1C and D). Consistent with these results, insulin sensitivity of Adipo-*Zip13*KO mice under HFD conditions was improved compared with that of Ctrl mice (Fig. S1E). Taken together, Adipo-*Zip13*KO mice appeared less susceptible to HFD induced insulin resistance.

**Fig. 1.**
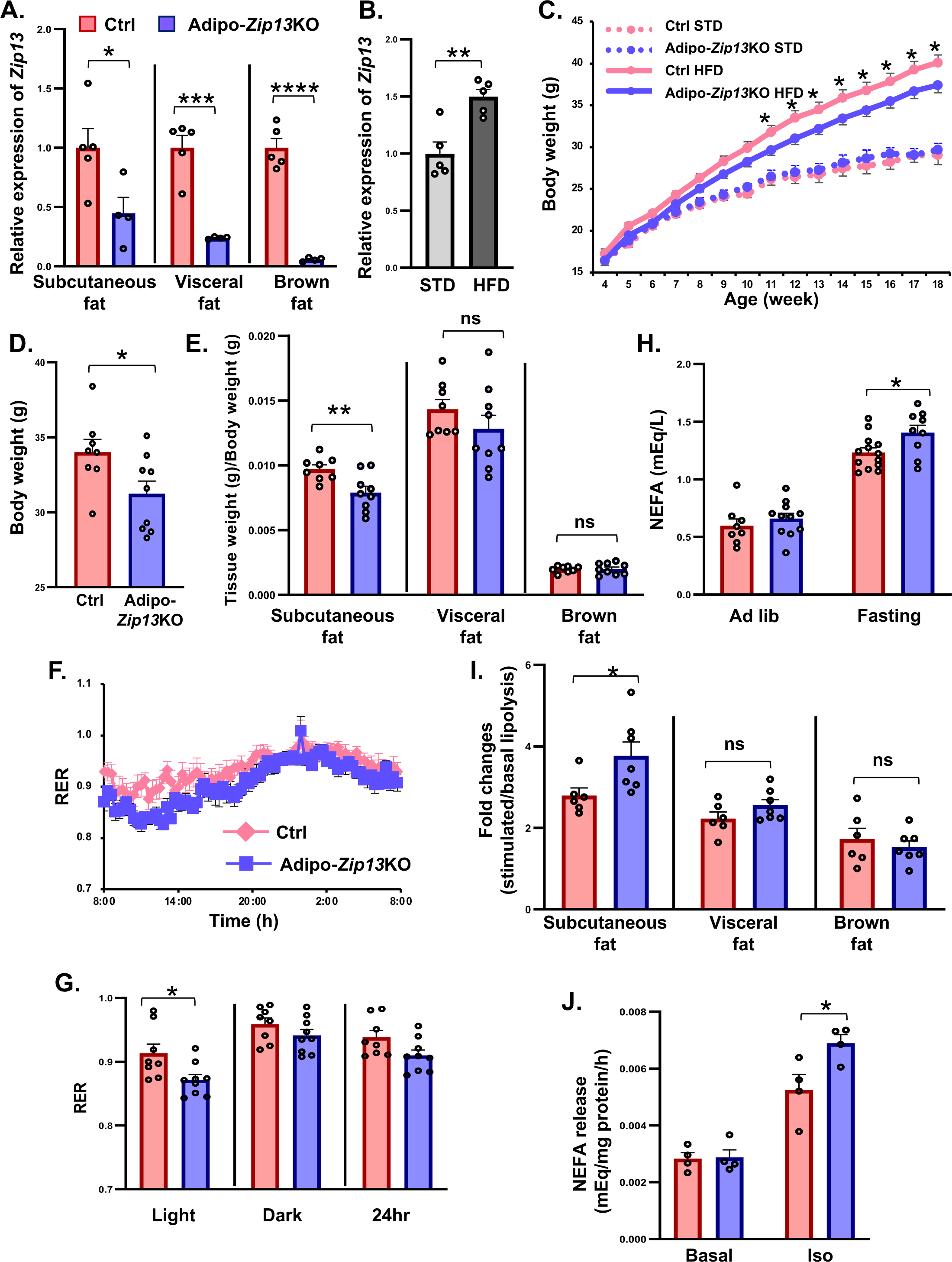
Adipo-*Zip13*KO mice accelerated lipolysis. (A) Relative mRNA levels of *Zip13* in the indicated tissues of 34-week-old Ctrl (n = 5) and Adipo-*Zip13*KO mice (n = 4). (B) Relative mRNA expression of *Zip13* in the subcutaneous fat tissue of WT mice (n = 5), fed STD or HFD for 12 weeks from 6-weeks old. (C) Body weights of the indicted mice fed a STD or HFD. Ctrl STD, n = 9 to 23; Adipo-*Zip13*KO STD, n = 7 to 19; Ctrl HFD, n = 17; Adipo-*Zip13*KO HFD, n = 22. (D) Body weight of 34-week-old Ctrl (n = 8) and Adipo-*Zip13*KO mice (n = 9). (E) Tissue weights of subcutaneous, visceral, and brown fat tissues of 34-week-old Ctrl (n = 8) and Adipo-*Zip13*KO (n = 9) mice. (F, G) Whole-body RER of 20-week-old Ctrl (n = 8) and Adipo-*Zip13*KO mice (n = 9) fed a STD. (F): RER trend, (G): RER period. (H) Plasma NEFA levels of 6-weeek old Ctrl (n = 8 to 13) and Adipo-*Zip13*KO mice (n = 9 to 11) fasted overnight and the i.p. injected with isoproterenol. (I) NEFA release from the indicated fat explants treated with saline (Basal) or isoproterenol (Iso). Fold changes are shown as stimulated/basal lipolysis. Analyzed fat pads were from 28 to 32-week old Ctrl (n = 6) and Adipo-*Zip13*KO mice (n = 7). (J) NEFA release from Ctrl and Adipo-*Zip13*KO differentiated cells treated with dimethyl sulfoxide (DMSO) (Basal) and isoproterenol (Iso). Data are shown as the mean ± SEM. (A, B, C, D, E, G, and I) \**p* < 0.05, \*\**p* < 0.01, by the two-tailed unpaired Student’s *t*-test. In (J and H), \**p* < 0.05, by two-way ANOVA followed by the Bonferroni’s multiple comparison test. ns, not significant.

To elucidate the mechanism of this phenotype observed in Adipo-*Zip13*KO mice, we investigated the metabolic state of the mice. Energy expenditure (Fig. S2A and B), food intake (Fig. S2C), locomotor activity (Fig. S2D), and rectal temperature (Fig. S2E) were comparable between Adipo-*Zip13*KO mice and Ctrl mice in STD condition. Gene expression analysis showed that modest but not statistically significant increases of *Ucp1* and *Cidea* expression are observed in the subcutaneous fat tissue of Adipo-*Zip13*KO mice (Fig. S2F). A similar gene expression pattern was observed in differentiated cells from subcutaneous fat tissue of Adipo-*Zip13*KO mice and these changes were also observed under the treatment with β-adrenergic receptor agonist isoproterenol (Fig. S2G). In addition, histology data showed the cell size of brown and white adipocytes is smaller in Adipo-*Zip13*KO mice under STD condition (Fig. S2H). We also measured the whole-body oxygen consumption (VO2) of Adipo-*Zip13*KO mice under different environmental temperatures. We found that their VO2 levels showed tendency to be increased (Fig. S2I), although the expression of brown fat markers was comparable (Fig. S2J). These results suggest that a modest increase in whole-body oxygen consumption may cause the resistance to high-fat diet-induced obesity and insulin resistance. Moreover, we found that RER was decreased in Adipo-*Zip13*KO mice (Fig. 1F and G: 20-week-old mice; Fig. S3A and B: 30-week-old mice in STD condition), suggesting that Adipo-*Zip13*-KO mice preferentially use lipids as an energy source rather than carbohydrates. We next investigated why Adipo-*Zip13*KO mice had decreased RER, then hypothesized that *Zip13*-deficient mature adipocytes might have accelerated lipolysis. To assess the degree of lipolysis, we measured circulating nonesterified fatty acid (NEFA) levels in the plasma of Adipo-*Zip13*KO mice in response to β-adrenergic stimulation, and found that their release of NEFA into the plasm was upregulated upon the administration of the isoproterenol, in the fasting condition (Fig. 1H), which suggests that the sympathetic nerve stimulated lipolysis was enhanced in Adipo-*Zip13*KO mice. We also found that subcutaneous fat tissue is the predominant fat tissue responsible for increased release of NEFA in Adipo-*Zip13*KO mice (Fig. 1I). We further investigated whether this accelerated lipolysis occurs in a cell-autonomous manner. We first isolated preadipocytes from the subcutaneous fat tissue of each mouse, and differentiated them into white adipocytes, followed by measurement of NEFA release into the medium. Unexpectedly, we found that NEFA release was increased in adipocytes of *Zip13*-deficient mice, upon isoproterenol stimulation in a cell-autonomous manner (Fig. 1J). Expression patterns of genes associated with lipolysis and lipid metabolism in the subcutaneous fat tissues of Adipo-*Zip13*KO mice were comparable with those of Ctrl mice (Fig. S3C). These results indicated that in Adipo-*Zip13*KO mice, lipolysis was accelerated in a cell-autonomous manner, even independent of the insulin level, without apparent changes in the expression levels of genes associated with the lipolysis process.

### Upregulation of cellular iron levels in mature adipocytes from Adipo-*Zip13*KO mice

We next aimed to investigate how ZIP13 is involved in the inhibition of lipolytic activity. It has been reported that mammalian ZIP13 transports zinc, whereas *Drosophil*a ZIP13 (dZIP13) transports iron ^23^, and therefore we assessed the cellular level of each metal using inductively coupled plasma mass spectrometry (ICP-MS) analysis and chemical probes (FluoZin-3 and FerroOrange for zinc ion (Zn^2+^) and the ferrous iron ion (Fe^2+^), respectively). Although zinc and iron levels in whole cells were not different between Ctrl and Adipo-*Zip13*KO mature adipocytes (Fig. 2A), organelle-associated Fe^2+^ levels, but not Zn^2+^ levels, were decreased in mature *Zip13* KO adipocytes (Fig. 2B). Based on reports that FerroOrange can label mainly endoplasmic reticulum (ER) and Golgi Fe^2+ 24^ and ZIP13 localized in both Golgi and ER (Fig. 2C), this result suggested that Fe^2+^ levels are decreased mainly in ER of mature *Zip13* KO adipocytes. We also analyzed the protein expression of iron regulatory protein 2 (IRP2), transferrin receptor (TfR), ferritin heavy chain (Fth1), and ferritin light chain (FTL), as the expression of these proteins is associated with iron metabolism. We found that the expression of both IRP2 and TfR was decreased by stimulation with isoproterenol, whereas the expression of Fth1 and FTL tended to be increased, although the Fth1 expression is not significantly (Fig. 2D and E). These results indicate that Fe^2+^ levels are increased in the cytosol of *Zip13* KO mature adipocytes, whereas they are decreased within their organelle including ER.

**Fig. 2.**
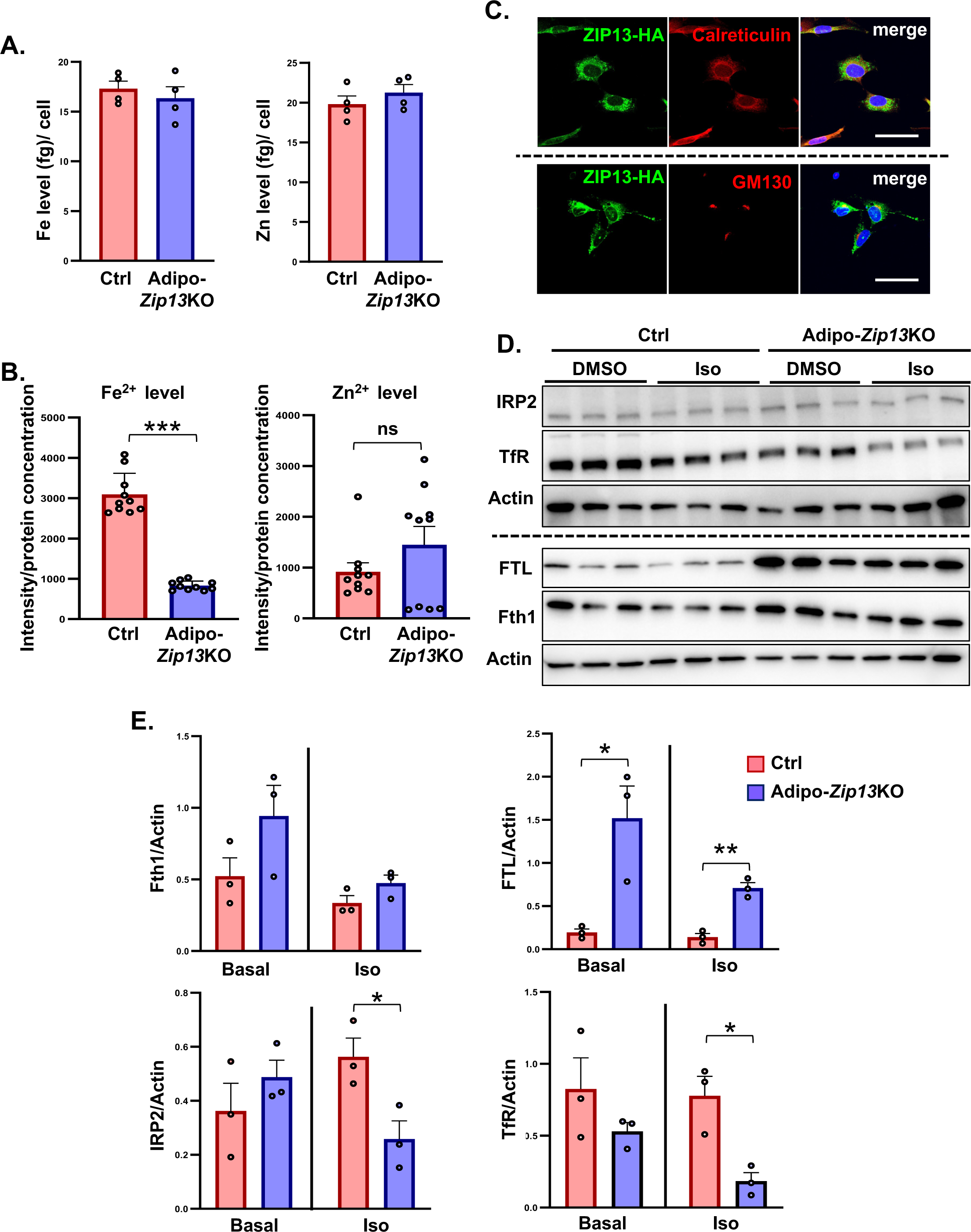
Adipo-*Zip13*KO mouse adipocytes increased intracytoplasmic iron levels but decreased organelles iron levels. (A) Zinc and iron levels in mature adipocytes. Zinc and iron level were measured by ICP-MS. (B) Zn^2+^ and Fe^2+^ levels in Ctrl and Adipo-*Zip13*KO mature adipocyte cells. Zn^2+^ and Fe^2+^ levels were detected by using FluoZin-3 and FerroOrange, respectively. (C) Immunofluorescence staining of mZIP13-HA in preadipocytes. Insertion of an HA sequence at C-terminal end allowed immunofluorescence studies using anti-HA antibody. Scale bar 50 μm. GM130: Golgi marker, Calreticulin, ER marker. (D) Immunoblotting of IRP2, TfR, FTL, Fth1, and actin using whole cell extracts of differentiated Ctrl and Adipo-*Zip13*KO cells treated with dimethyl sulfoxide (DMSO) (Basal) or isoproterenol (Iso). Actin was included as a loading control. (E) Each protein shown in (D) was quantified by normalization to actin. Data are shown as the mean ± SEM. In (A, B and E), \**p* < 0.05, \*\**p* < 0.01, \*\*\**p* < 0.001, by the two-tailed unpaired Student’s *t*-test.

### ZIP13-mediated iron negatively regulates lipolytic activity

Next, we assessed whether the ectopic expression of ZIP13 could increase the Fe^2+^ levels of organelles. Either the wild-type form of mouse ZIP13 (ZIP13-WT) or its H254A mutant (ZIP13-H254A), in which metal transport activity is attenuated ^18, 25^, was retrovirally expressed in *Zip13* KO preadipocytes, followed by their differentiation into mature adipocytes. The cells were then harvested on the second day (referred to as preadipocytes) or the sixth day (referred to as mature adipocytes) (Fig. 3A). In the preadipocytes, we observed that the expression of ZIP13-WT induced an increase in cellular Zn^2+^ but not Fe^2+^ levels in organelles, and this change was reduced in the expression of ZIP13-H254A compared to ZIP13-WT expression (Fig. 3B). Intriguingly, in mature adipocytes, not only cellular Zn^2+^ but also Fe^2+^ levels in organelles were increased by ZIP13-WT, although only Fe^2+^ levels in organelles were reduced by the expression of ZIP13-H254A compared to ZIP13-WT expression (Fig. 3C). These results suggest that ZIP13 may be involved in cellular zinc and iron homeostasis in different ways depending on the adipocyte stage.

**Fig. 3.**
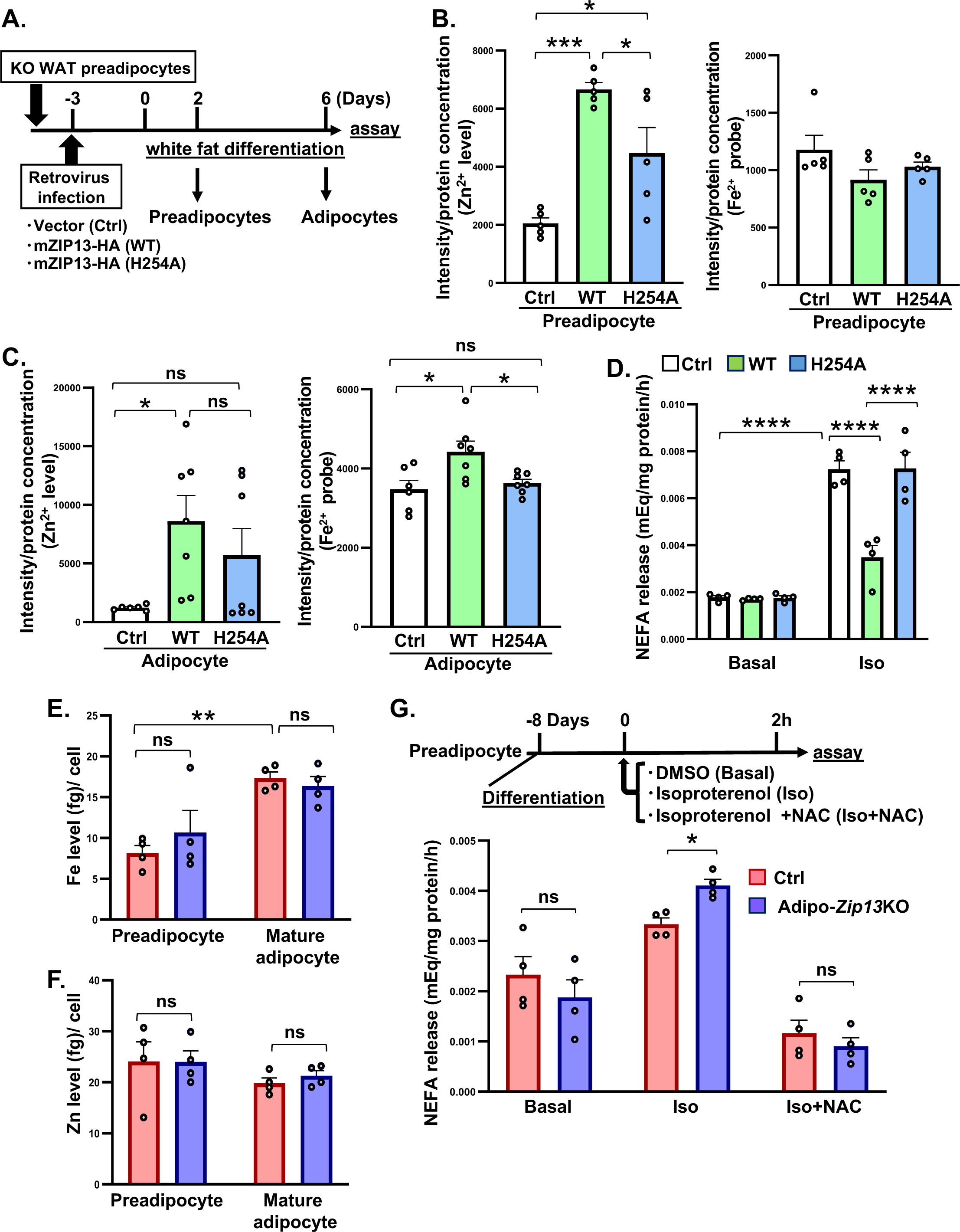
ZIP13-mediated iron negatively regulates lipolysis. (A) Schematic representation of the experimental plan. (B) Zn^2+^ and Fe^2+^ level in *Zip13* KO preadipocyte cells expressing ZIP13-WT and ZIP13-H254A. (C) Zn^2+^ and Fe^2+^ levels in *Zip13* KO mature adipocyte cells expressing ZIP13-WT and ZIP13-H254A. (D) NEFA release from Adipo-*Zip13*KO mature adipocyte cells expressing ZIP13-WT or ZIP13-H254A, and treated with DMSO (Basal) and isoproterenol (Iso). (E) Total iron levels of Ctrl and Adipo-*Zip13*KO preadipocyte and mature adipocyte cells. (F) Total zinc levels of Ctrl and Adipo-*Zip13*KO preadipocyte and mature adipocyte cells. (G) Schematic representation of the experimental plan (upper panel). Ctrl and Adipo-*Zip13*KO cells were differentiated by stimulation with isoproterenol, in the presence or absence of N-acetylcysteine (NAC), and then NEFA release was analyzed (lower panel). Data are shown as the mean ± SEM. In (B, C, D, E, and F), \**p* < 0.05, \*\**p* < 0.01, \*\*\**p* < 0.001, \*\*\*\**p* < 0.0001, by one-way ANOVA followed by the post-hoc Tukey-Kramer test. In (G), \**p* < 0.05, by the two-tailed unpaired Student’s *t*-test. ns, not significant.

We next aimed to determine whether ZIP13-mediated metal transport is involved in the lipolysis of mature adipocytes, using the mutant ZIP13-H254A, which impaired Fe^2+^ transporting ability in mature adipocytes (Fig. 3C). We found that ZIP13-H254A could not suppress NEFA release like ZIP13-WT does (Fig. 3D), suggesting that the Fe^2+^ transport ability of ZIP13 in the mature adipocyte is required for the inhibition of lipolysis. Notably, ICP-MS analysis indicated that amounts of iron were significantly increased in mature adipocytes, whereas their zinc levels were comparable or rather tended to be lower than that of preadipocytes (Fig. 3E and F). These results suggest that ZIP13 can transport both Zn^2+^ and Fe^2+^, and that ZIP13-mediated Fe^2+^ plays a crucial role in negatively regulating lipolysis in mature adipocytes. This result was reminiscent of a previous study showing that iron-mediated lipolysis is caused by a pro-oxidant mechanism, in which supplementation of the anti-oxidant N-acetyl cysteine (NAC) was shown to cancel iron-induced NEFA release ^26^. Therefore, we tested whether NAC treatment can reduce lipolysis in *Zip13*-deficient mature adipocytes, and NAC treatment significantly inhibits NEFA release in mature adipocytes (Fig. 3G). Furthermore, hydroxy-radicals produced by Fe^2+^ mediated the Fenton reaction was increased in Adipo-*Zip13*KO cells (Fig. S4), indicating that ZIP13-mediated Fe^2+^ sequestration in ER may be involved in the degree of lipolysis, and dysfunction of ZIP13 increases cytoplasmic Fe^2+^ and pro-oxidants levels, leading to NEFA release.

### Phosphodiesterase (PDE) activity is reduced in Adipo-*Zip13*KO cells

How does ZIP13-mediated Fe^2+^ transport affect the lipolysis process? To address this question, we focused on the β-adrenergic signaling pathway, which activates the lipolysis process in a cyclic AMP (cAMP) and protein kinase A (PKA)-dependent manner ^27^. We found that both levels of cAMP and phosphorylated PKA (pPKA) were increased in *Zip13* KO mature adipocytes (Fig. 4A and B, Fig. S5), suggesting that the steps upstream of cAMP are activated in these cells. On the other hand, NEFA release from *Zip13*KO mature adipocytes was increased even upon treatment with forskolin, an activator of adenylate cyclase (Fig. 4C), suggesting that the steps downstream of adenylate cyclase would be affected in *Zip13* KO mature adipocytes. Then, we assessed the activity of PDE, which degrades cAMP, and found that its activity was decreased in *Zip13* KO mature adipocytes (Fig. 4D), suggesting that the decrease in the PDE activity would be responsible for the acceleration of lipolysis by the loss of ZIP13. To confirm this point, we focused on PDE3 and PDE4, which are both predominantly expressed in adipose tissues ^28^, and treated the cells with each inhibitor to see which PDE predominantly plays a role in lipolysis. We found that the inhibition of PDE3 increased NEFA release from Ctrl mature adipocytes but not from *Zip13* KO mature adipocytes (Fig. 4E), and such increased NEFA release was not observed using a PDE4 inhibitor, suggesting that perturbation of PDE3 may occur in *Zip13*-deficient mature adipocytes, which might be the main reason for the increased cAMP level leading to the acceleration of lipolysis.

**Fig. 4.**
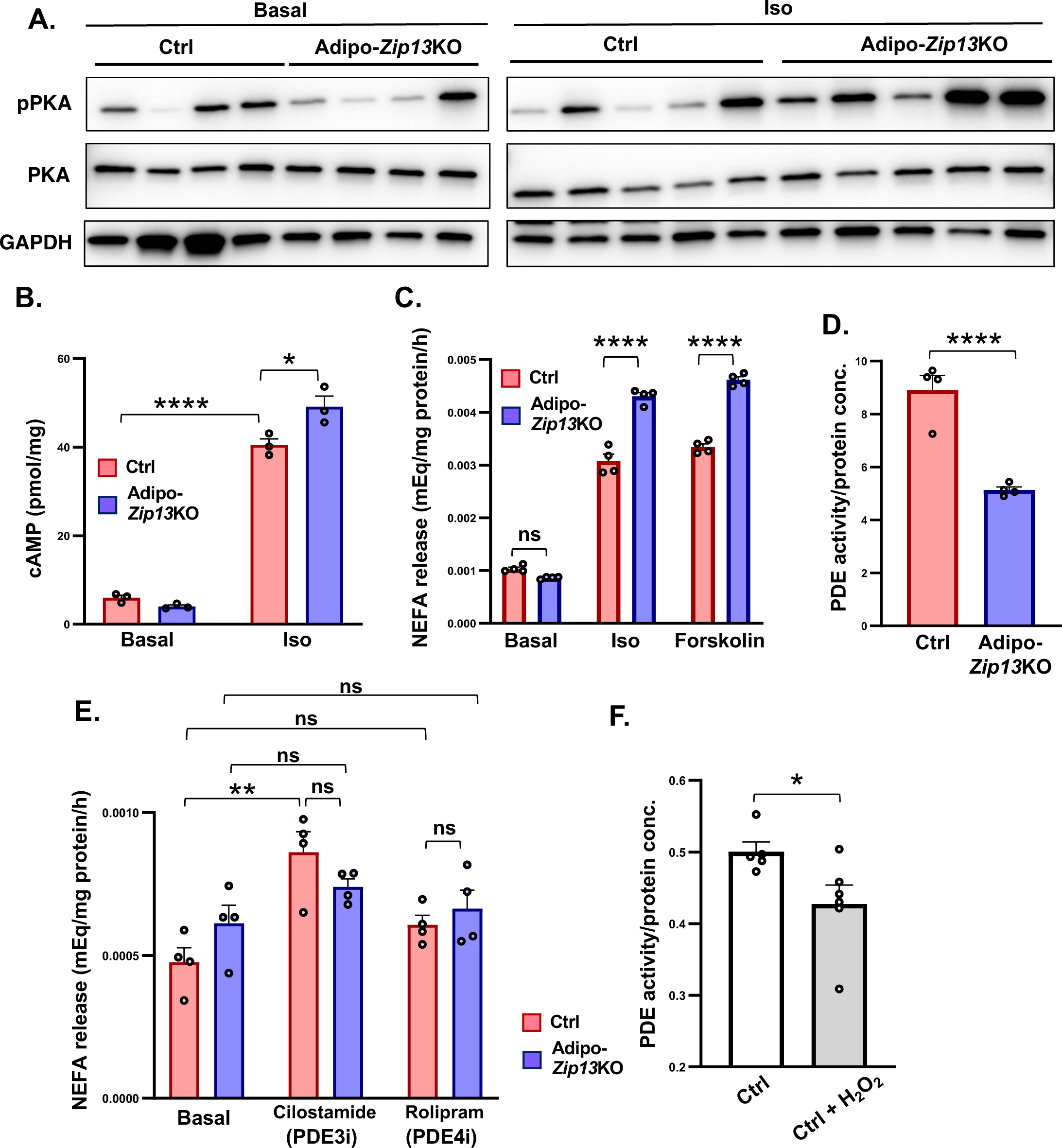
PDE activity is reduced in Adipo-*Zip13*KO cells. (A) Immunoblotting of pPKA, PKA, and GAPDH in subcutaneous fat tissues from Ctrl and Adipo-*Zip13*KO mice, stimulated with DMSO (basal) (left panel) or isoproterenol (Iso) (right panel). (B) cAMP levels of Ctrl and Adipo-*Zip13*KO mature adipocytes in the presence (Iso) or absence (Basal) of isoproterenol. (C) NEFA release from differentiated Ctrl and Adipo-*Zip13*KO mature adipocytes stimulated with isoproterenol, or forskolin. (D) PDE activity in Ctrl and Adipo-*Zip13*KO mature adipocytes, normalized by protein concentration. (E) NEFA release from the differentiated Ctrl and Adipo-*Zip13*KO cells, stimulated with cilostamide (PDE3 inhibitor), or rolipram (PDE4 inhibitor). (F) PDE activity normalized by protein concentration of Ctrl adipocyte cells in the presence or absence of H_2_O_2_. Data are shown as the mean ± SEM. In (A, D, and F), \**p* < 0.05, \*\*\*\**p* < 0.0001, by the two-tailed unpaired Student’s *t*-test. In (B and C), \**p* < 0.05, \*\*\*\**p* < 0.0001, by one-way ANOVA followed by the post-hoc Tukey-Kramer test. In (E), \*\**p* < 0.01, by two-way ANOVA followed by the Bonferroni’s multiple comparison test. ns, not significant.

As *Zip13*-deficient mature adipocytes have increased Fe^2+^ in the cytosol, and pro-oxidants are involved in facilitating lipolysis ^26^, we next investigated whether oxidative stress affects PDE activity. We demonstrated that PDE activity was decreased by H_2_O_2_ treatment (Fig. 4F), which was consistent with a previous report ^29^. These results suggest that *Zip13*-deficient mature adipocytes have increased Fe^2+^ in the cytosol and increased pro-oxidants, leading to the inhibition of PDE3, which results in the enhancement of NEFA release.

### Mammalian ZIP13 transports both iron and zinc ions

Crucial questions to be clarified were how *Zip13* deficiency caused the changes in the subcellular distribution of Fe^2+^ level in mature adipocytes, and whether or not mammalian ZIP13 could transport iron directly, as ZIP13 has been recognized as a zinc-specific transporter that regulates intracellular zinc homeostasis ^15,16, 17, 18^. Although some ZIP family members, such as ZIP8 and ZIP14, transport multiple metal ions, including zinc and iron ^10^, direct evidence of the transportation of Fe^2+^ by mammalian ZIP13 has not been reported to date. To address these questions, we heterologously expressed ZIP13 on the plasma membrane of *Xenopus* oocyte and examined its transport activity of zinc and iron ions.. Upon injection of complementary RNA (cRNA) of the full-length form of mouse ZIP13 (mZIP13), its protein was mainly expressed in intracellular spaces but not on the cell surface of oocytes (Fig. 5A, upper left). This is the issue that has made it difficult to experimentally determine substrates of ZIP13 that is transported, as ZIP13 is an intracellular transporter that localizes to the ER/Golgi in mammalian cells. To circumvent this problem, we fused mZIP13 with human SLC3A24/4F2 cell-surface antigen heavy chain (4F2hc), which is the subunit of this large neutral amino acid transporters such as LAT1, to force the ZIP13 protein to be localized on the plasma membrane (Fig. 5B). We confirmed that this fusion protein named 4F2hc-mZIP13-HA (4F2hc-m13) was successfully and predominantly expressed on the surface of oocytes (Fig. 5A, lower left), which was also true for the case of another fusion protein, 4F2hc-mZIP14-HA (4F2hc-m14), which was used as a control (Fig. 5A, lower right). Three-dimensional structure simulation using Alphafold2 suggested that the structure of the metal transporting regions of these proteins might be unaffected (Fig. 5C), and both 4F2hc-m13 and 4F2hc-m14 induced the expression of metallothionein 1 gene *(MT1)*, an indicator of intracellular zinc level ^16, 17, 18^ in HEK293 cells when each of them was ectopically expressed (Fig. 5D), indicating that both fusion transporters can transport zinc into cells. Then, we assessed if these fusion transporters expressed on the plasma membrane of *Xenopus oocytes* are able to mobilize zinc and iron ions, using the *Xenopus* oocyte transport assay with radioisotopes, and demonstrated that the ZIP14-based fusion transporter 4F2hc-m14 as well as the ZIP13-based fusion transporter 4F2hc-m13 transported zinc and iron (Fig. 5E and F). These results clearly demonstrated that mammalian ZIP13 has the ability to transport both zinc and iron ions.

**Fig. 5.**
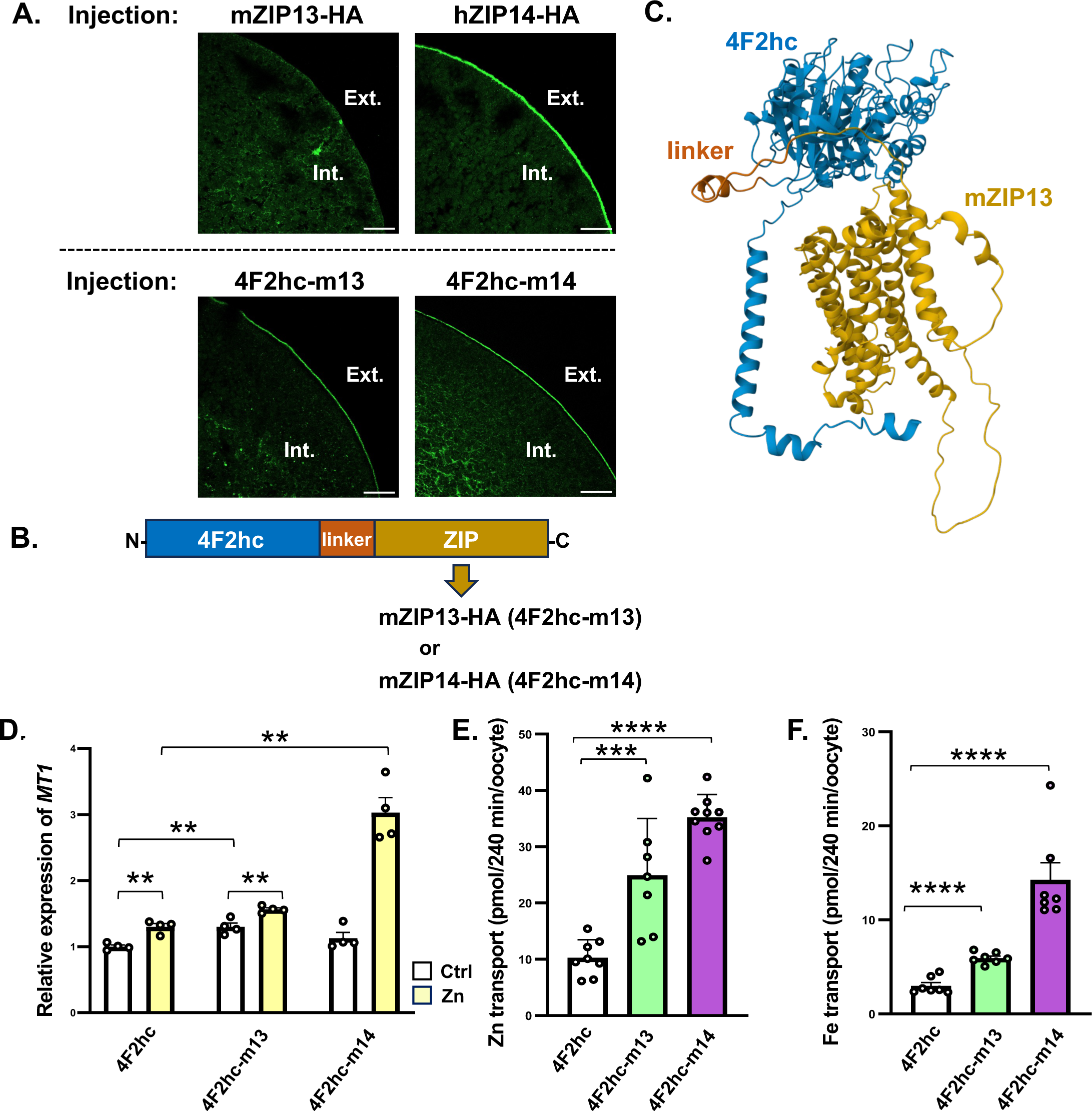
ZIP13 has the ability to transport both zinc and iron. (A) Immunofluorescence microscopic detection of the expression of mZIP13, hZIP14, 4F2hc-m13, and 4F2hc-m14 in *X*. *laevis* oocytes. Insertion of an HA sequence at the C-terminal end enabled immunofluorescence analyses using an anti-HA antibody. Scale bar = 50 μm. (B) Schematic illustration of 4F2hc-m13 and 4F2hc-m14. (C) Alphafold2 simulation of the fusion protein of 4F2hc and mZIP13. (D) Relative *MT1* expression in HEK293 cells expressing 4F2hc, 4F2hc-m13, or 4F2hc-m14 with the addition of 20 μM ZnSO_4_ was added. (E) Zinc uptake of 4F2hc-m13 and 4F2hc-m14. (F) Iron uptake of 4F2hc-m13 and 4F2hc-m14. Data are shown as the mean ± SEM. In (C), \**p* < 0.05, \*\**p* < 0.01, \*\*\**p* < 0.001, \*\*\*\**p* < 0.0001, by one-way ANOVA followed by the post-hoc Tukey-Kramer test. In (E and F), \*\*\*\**p* < 0.0001, by the two-tailed unpaired Student’s *t*-test. Ext. extracellular space, Int. intracellular space.

### *In silico* structural dynamics simulations of transport mechanism of mammalian ZIP13

Lastly, we performed an *in silico* ion binding prediction in terms of static and dynamic analyses using the ZincBind web server and molecular dynamics (MD) simulation, respectively. The result predicted that Zn^2+^ could bind to M1 and M2 sites, which is relevant to N120, D225, N226, H229, H254, E255, and E259 (Fig. 6A), similar to what was previously reported in ZIP8 ^30^. The ion binding stability of Zn^2+^ and Fe^2+^ was further evaluated using 0.5 μs-MD simulations for the wild-type form of mouse ZIP13 (ZIP13-WT), with 3 replications for each system, totaling 6 systems. The results showed a consistent trend across all replications of each system. However, it was observed that only in #3, Zn^2+^ was trapped by H124 at the M1 site over the simulation time. For the representative systems, this ion-binding analysis shows that both Zn^2+^ and Fe^2+^ at the M2 site could stably bind to ZIP13-WT/Zn^2+^ and ZIP13-WT/Fe^2+^ after 0.33 μs (Fig. 6A and B). In contrast, ions at the M1 site slightly fluctuated and subsequently moved out of this ion channel, particularly moving out faster in the ZIP13-WT/Fe^2+^ compared to the ZIP13-WT/Zn^2+^ (Fig. 6A and B). Notably, Zn^2+^ at the M1 site in ZIP13-WT stably bound until 0.17 μs, while the ion at M1 site of ZIP13-WT/Fe^2+^ system rapidly moved out around 0.05 μs in all replications (Fig. 6B). Based on three replicates of each system, we found that in one of the ZIP13-WT/Zn^2+^ simulations (run #3), Zn^2+^ at M1 remained bound throughout the simulation time. This might indicate the superiority in Fe^2+^ transport kinetics. Among amino acid residues in the M1 and M2 sites, E255 and E259 show a strong binding interaction with both ions according to the occupancy of the ion at each residue (Fig. 6C). When the ion is released from the M1 site, conformational changes in the side chain of H254 facilitate ion ejection, confirming the important role of H254 in the ion transport of ZIP13. In this process, H254 binds Zn^2+^ slightly longer than Fe^2+^, which is probably the reason why Fe^2+^ is ejected faster from the ion binding sites. These simulation results also support our experimental results that ZIP13 may serve a role in transporting both Zn^2+^ and Fe^2+^ ions.

**Fig. 6.**
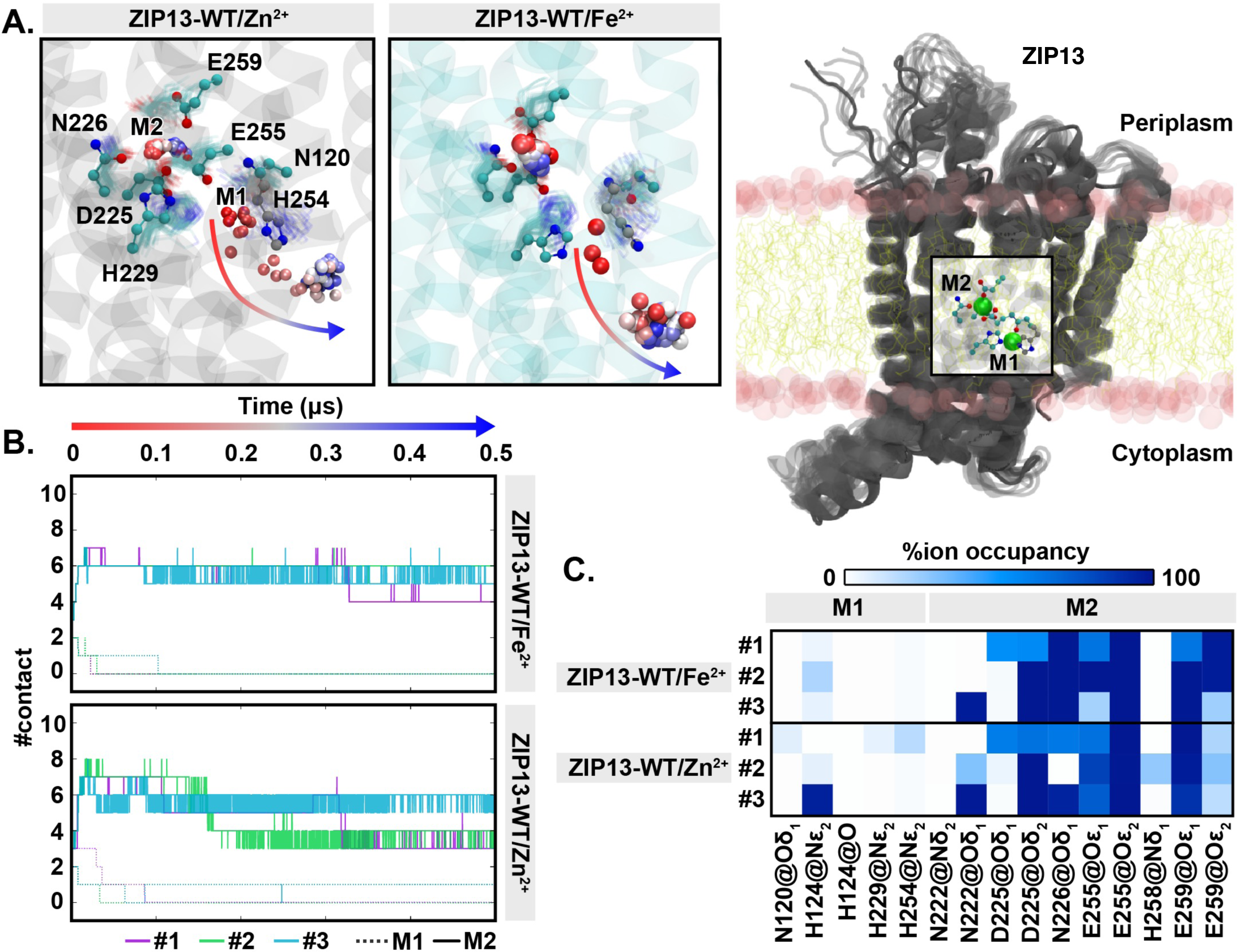
Ion binding site prediction and analysis of mouse ZIP13 protein. (A) The ion binding site of representative structure of ZIP13 at M1 and M2 embedded in lipid membrane (right panel) and motion of ions and amino acid residues in the M1 and M2 sites, where red to blue color change indicates the time lapse (left panel). (B) The number of atom contacts (#contact) to ion during the 0.5 μs-MD simulations. (C) Occupancy of Zn^2+^/Fe^2+^ ions nearby amino acid residues at the M1 and M2 sites. Oδ and Oε mean two different oxygens in the carboxy group, and Nε does the Nε atom in the histidine.

## Discussion

Dysregulation of iron homeostasis is associated with several disease statuses including diabetes and obesity ^31^. For instance, high ferritin levels are positively correlated with central adiposity ^32^, and iron deficiency is also associated with obesity ^33, 34^. It has been reported that iron regulates adipocyte function and homeostasis associated with adipogenesis and lipid metabolism. It is widely known that iron is crucial for the generation of lipid peroxidation products, which are linked with many pathophysiological statuses, and it is also true that iron overload under the condition of obesity increases lipolysis and lipid peroxidation ^26^. Although the role of iron in lipid metabolism has been well-recognized, the molecular mechanism of how iron regulates lipid metabolism in adipose tissues *in vivo* and its physiological significance is still poorly understood.

As we previously found that ZIP13 participates in adipose tissue homeostasis by regulating the adipocyte browning ^18^, we herein investigated how ZIP13 is involved in maturing adipocyte functions. We found that total iron and zinc levels in cells were comparable between control and *Zip13* KO mature adipocytes (Fig. 2A), however, Fe^2+^ levels in the cytosol of *Zip13* KO mature adipocytes were significantly increased (Fig. 2D and E) and correlated with the promotion of lipolysis (Fig. 3). We also found that the cAMP/PKA signaling pathway was upregulated in *Zip13* KO mature adipocytes, which was owing to the downregulation of PDE activity (most likely of PDE3, Fig. 4E). This inhibition of PDE activity is due to increased reactive oxygen species (ROS), as the ROS scavenger NAC moderates NEFA release. These results indicate that enhancement of the intracellular iron-ROS axis in the absence of ZIP13 attenuates PDE activity in mature adipocytes, leading to an increase in cAMP level and lipolysis (Fig. 7). To our knowledge, it was the first to demonstrate that mammalian ZIP13 affects subcellular iron distribution without changing total cellular iron levels. Although, the gross phenotypes of *Zip13* KO mice are reminiscent of those of the zinc-deficient model suffering short stature and skin abnormalities, which are different from the phenotypes of iron deficiency. On the other hand, it is also known that all the phenotypes or symptoms of *Zip13* KO or EDSSPD3 patients cannot be explained solely by zinc dysregulation. Therefore, the ZIP13 mediated intracellular transport of iron may be a novel mechanism that contributes to the lipolysis in mature adipocytes *in vitro* and *in vivo*.

**Fig. 7.**
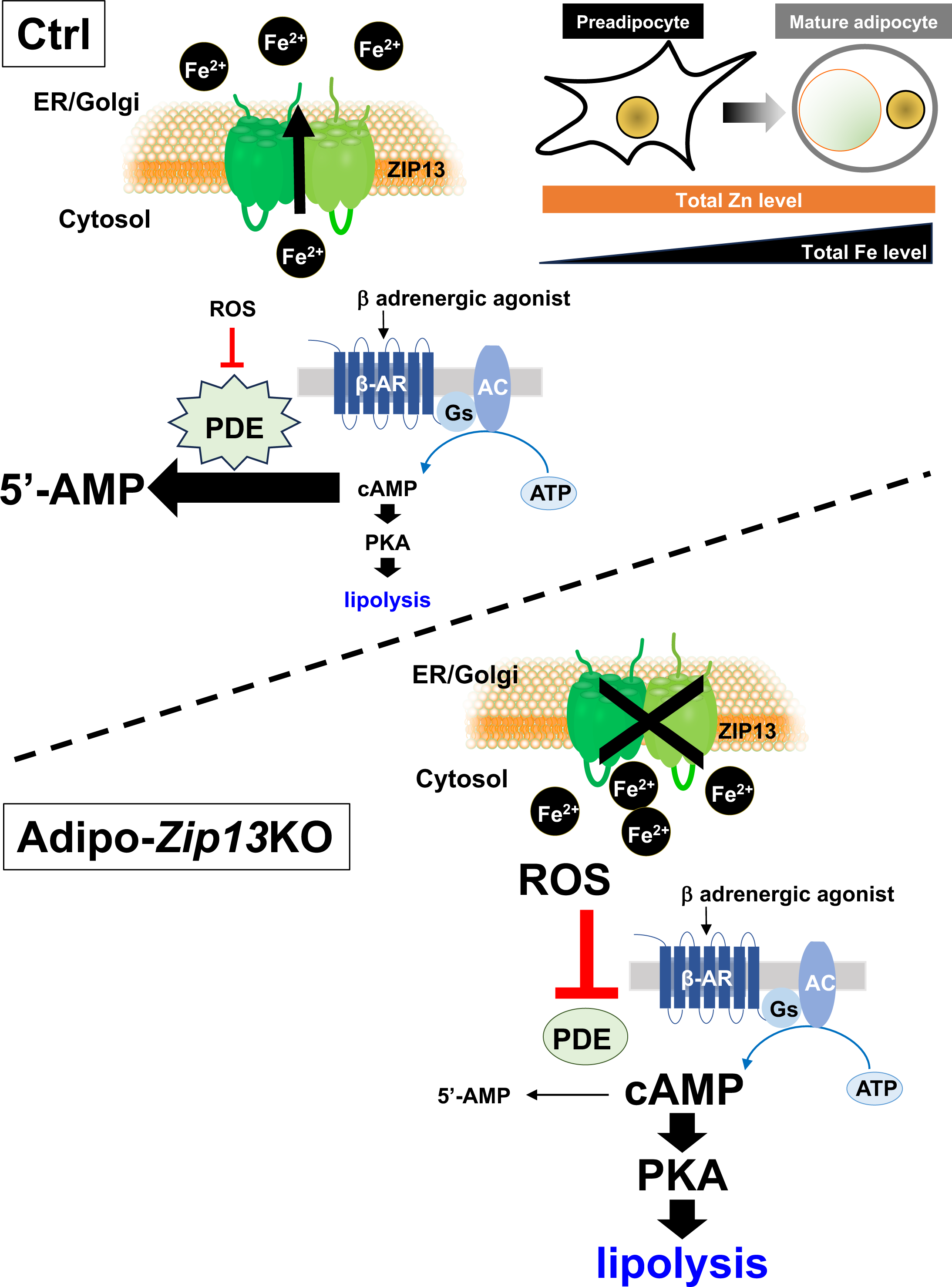
ZIP13 regulates adipocyte lipolysis through ZIP13-mediated iron. Schematic illustration of metal ion balance in preadipocytes and mature adipocytes. Zinc level is slightly decreased during adipocyte differentiation, whereas iron level is dramatically increased during adipocyte differentiation (upper right panel). ZIP13-mediated iron suppresses the β-adrenergic receptor signaling pathway, thereby inhibiting lipolysis.

EDSSPD3 patients were found to have lipoatrophy and decreased adipose tissue mass, which are closely associated with the phenotypes observed in Adipo-*Zip13*KO mice. A recent study showed that there is an adipogenic progenitor cell population that expresses *Acta2* (encoding smooth muscle actin (αSMA), and *Sm22*, *Pax3*, *Pdgfrα,* and *Cd81* were identified in the stromal vascular fraction (SVF) of white adipose tissue (WAT) as precursors of beige adipocyte following cold exposure ^35, 36, 37, 38^. Furthermore, WAT lipolysis-derived linoleic acid triggers beige progenitor cell proliferation following cold acclimation, and β3-adorenoceptor activation ^39^. As the expression of *Zip13* was also observed in the SVF in addition to the adipocyte fraction (Fig. S6A), and CD81-positive proliferating cells were increased in the SVF of *Zip13* KO mice (Fig. S6B), there are possibilities that ZIP13-mediated lipolysis in mature adipocyte might be involved in the proliferation of beige progenitor cell or that ZIP13 might be directly involved in the regulation of beige progenitor cells. Further analysis is needed to elucidate the role of ZIP13 in beige progenitor cells and/or beige fat biogenesis.

We previously demonstrated that ZIP13-mediated zinc is necessary for the inhibition of beige adipogenesis ^18^. In the present study, we found that ZIP13 has the ability to transport both Fe^2+^ and Zn^2+^. Furthermore, we also found that H254 residue of ZIP13 is important for the transport Fe^2+^ and Zn^2+^ by using both cell experiments (Fig. 3B and C) and *in silico* structural dynamics simulations (Fig. 6), and that ZIP13-mediated iron distribution in mature adipocytes is important for proper lipolytic function (Fig. 3D). Although it is well recognized that iron is an important metal affecting many biological functions, its significance in subcellular distribution has remained poorly characterized to date. Our results demonstrated that iron balance between cytosol and intracellular vesicles such as ER and Golgi achieved through ZIP13 affect the degree of lipolysis induced by β-adrenergic stimulation in the mature adipocyte. It is still unknown whether ZIP13 is the only molecule that affect intracellular sequestration of iron ion, these results points to future investigation into the elucidation of regulatory mechanism of intracellular iron distribution.

It is also an intriguing finding that ZIP13 may act as a dual-specificity transporter of Zn^2+^ and Fe^2+^ during the maturation of adipocytes. In a metal ion transport assay using *E.coli*., dZIP13 was found to transport only Fe^2+^, unlike mammalian ZIP13 ^23^. Furthermore, in contrast to mammalian ZIP13, dZIP13 has potential iron-binding residues at both its N- and C-termini ^40, 41^. This is consistent with the fact that dZIP13 responds to iron and plays a specific role in iron homeostasis. In contrast, mZIP13 regulates only zinc levels in preadipocyte cells, whereas mZIP13 regulates both iron and zinc levels in mature adipocytes in our adipocyte differentiation system (Fig. 2 and Fig. 3), suggesting that mZIP13 transports both Fe^2+^ and Zn^2+^ within cells in a context-dependent manner. The question then arises as to how the specificity of substrate transport is determined in cells and tissues. One possible factor that determines substrate specificity may be changes in intracellular and extracellular environment. Notably, several studies reported that iron levels are increased during adipocyte differentiation ^42, 43^. Based on our (Fig. 3E and F) and other observations ^42, 43^, it is possible that mZIP13 switches the main physiological substrate from Zn^2+^ to Fe^2+^ during adipocyte differentiation, possibly responding to an increase in intracellular iron concentrations. To investigate our hypothesis, we performed *in silico* structural dynamics simulation based on actual zinc and iron levels measured by ICP-MS, and found that mZIP13 may preferentially transport Fe^2+^ over Zn^2+^ (Fig. 6). Further analyses are needed to clarify the signals or timing that is important for “possible switching from zinc to iron transport” by ZIP13.

Of particular importance in this study is the establishment of a transport assay system for ZIP family existing on the membrane of intracellular compartment, such as ER or Golgi. Using 4F2hc fusion ZIP transporters, we successfully expressed ZIP fusion proteins on the plasma membrane of *Xenopus oocytes* and examine zinc and iron ion transport in the system (Fig. 5). Recent advances in cryo-electron microscopy showed that, BbZIP, a ZIP homolog in *Bordetella bronchiseptica*, transports Zn^2+^ in an elevator-type mechanism ^44, 45, 46^; however, the molecular mechanism of ion transport by ZIPs has not yet been clarified, although Zn^2+^/HCO^3-^ and Zn^2+^/H^+^ symport or water-mediated transport mechanisms have been proposed ^47, 48^. We believe that clarification of detailed transport mechanisms of zinc and iron ions by ZIPs is the important next step in the field, for which the established transport assay system in this study will contribute in creating a venue that highlights and explores the various facets of ZIP family members.

In conclusion, our findings illustrate that ZIP13 acts as a key regulator for lipolysis in adipocytes via the proper use of metals, and a picture is now emerging of the ZIP13-iron axis plays an important role in regulation of lipid metabolism. With this realization that ZIP13 is a critical regulator of intracellular metal homeostasis and lipid metabolism, it may become a focus of attention for its potential to identify novel targets for therapeutic intervention in human disease.

## Methods

### Animal studies

All mice were housed in specific pathogen-free barrier facilities, maintained under a 12-h light/dark cycle, given water *ad libitum*, and fed standard rodent food (Oriental Yeast, Tokyo, Japan) or rodent food containing 60% fat (Research Diet, New Brunswick, NJ, USA) from 6 to 18 weeks of age ^49^. *Zip13*-flox mice (Accession No. CDB1360K: https://large.riken.jp/distribution/mutant-list.html) were generated by using TT2 embryonic stem cells derived from a hybrid strain between C57BL/6 and CBA ^50^. To construct a targeting vector, genomic fragments of the *Zip13* locus were obtained from a C57BL/6 BAC clone (BACPAC Resources). A 3.8 kbp region containing from exon 5 to 10 of the *Zip13* gene was flanked by loxP sites (Fig. S7). The targeted ES clones were microinjected into 8-cell stage ICR embryos, and the injected embryos were transferred into pseudopregnant ICR females. The resulting chimeras were crossed with B6;SJL-Tg(ACTFLPe)9205Dym/J mice to remove the drug-resistant Neo and Puro cassettes, and heterozygous offspring were identified by PCR using the following primers: P1 (5’-CCA GCC AGGT GAG CCC CAA G-3’) and P2 (5’-TTC CGG GCA GAG GGC ACA GT-3’) (WT; 1449 bp, Flox; 1730 bp), and P3 (5’-ACCGGCTGCCTCTTCCACAT-3’) and P4 (5’-GTG CCC CCT GCT GTG TGA GG-3’) (WT; 513 bp, Flox; 689 bp). Adiponectin-Cre mice were obtained from Jackson Laboratories (strain no. 010803), and all mice were backcrossed onto C57BL/6J mice for more than seven generations. Adipocyte-specific *Zip13* KO mice were generated by crossing *Zip13-*flox mice with adiponectin-Cre mice. *Zip13f/f;* adiponectin-Cre mice were initially used as adipocyte-specific *Zip13* KO mice, however, as they showed no obvious phenotypes associated with mature adipocytes such as RER and VO2 values as observed in systemic *Zip13*KO mice, then we used *Zip13f/*(−);adiponectin-Cre mice were generated and used, because of their higher efficiency of KO compared with *Zip13f/f;* adiponectin-Cre mice. In this study, we used *Zip13f/*(−); adiponectin-Cre mice were used as Adipo-*Zip13*KO mice, and *Zip13f/*(+); adiponectin-Cre mice were used as Ctrl mice.

### Metabolic studies Indirect calorimetry

VO2 and VCO2 were measured in individual mice at the indicated ages using an Oxymax apparatus (Columbus Instruments, Columbus, OH, USA). O_2_ and CO_2_ measurements were performed every 18 min over a 3-day period, during both the light and dark phases, and the data from the final day were analyzed. Locomotor activity was measured in a metabolic chamber using an infrared light beam detection system (ACTIMO-100; Shinfactory, Fukuoka, Japan). Total locomotor activity during a 12-h period was measured.

### Cell immortalization and culture

White preadipocytes were isolated from the subcutaneous fat tissue of Ctrl and Adipo-*Zip13*KO mice (6 -8 weeks) by collagenase digestion, and then immortalized, as described previously ^18^. Briefly, preadipocytes were immortalized by retroviral transduction with the SV40 largeT antigens, and selection with puromycin (2 mg/mL). Immortalized preadipocytes were a mixed population, and two cell lines were analyzed. Preadipocytes were seeded onto collagen-coated dishes (Corning, Kennebunk, ME, USA) in Dulbecco’s Modified Eagle Medium (DMEM)/F12 (Gibco, Carlsbad, CA, USA) with 10% fetal bovine serum (FBS). For white adipocyte differentiation, cells were induced with induction medium containing 10% FBS, 5 μg/mL insulin (Humalin R, Eli Lilly), 250 μM isobutylmethylxanthine (Sigma-Aldrich, St. Louis, MO, USA), 125 μM Indomethacin (Sigma), and 2 μg/mL dexamethasone (Sigma) in DMEM/F12 according to the manufacturer’s protocol ^51^. Two days after induction, the culture medium was changed to a maintenance medium containing 10% FBS and 5 μg/mL insulin. For isoproterenol treatment, cells were incubated with 10 μM isoproterenol for 2 h.

### Immunoblotting experiments

Immunoblotting was performed as described previously ^18, 52, 53, 54^. The following antibodies were used for immunoblotting: anti-Fth1 (1:1,000; Abcam, St. Charles, MO, #EPR3004Y), anti-FTL (1:1,000; Abcam, #ab69090), anti-IRP2 (1:1,000; Millipore, Bedford, MA, USA, #MABS2030), anti-TfR (1:1000; Invitrogen, Carlsbad, CA, USA, #136890), anti-actin (1:5000, Sigma, #A2228), anti-pPKA (1:1000; Cell Signaling, Danvers, MA #5661), PKA (1:1000; Cell Signaling, #5842) and anti-GAPDH (1:3,000; Cell Signaling, #2118).

### Immunofluorescence experiment

*Xenopu*s *laevis* oocytes injected with cRNAs were fixed with paraformaldehyde and embedded in paraffin. The sliced sections were incubated with an anti-HA monoclonal antibody (1:1000; BioLegend, San Diego, CA, USA, #901514), followed by anti-mouse Alex Fluor 488-conjugated immunoglobulin (1:100; Thermo Fisher Scientific, Tokyo, Japan). The sections were mounted in Fluorescence mounting medium (DAKO, Glostrup, Denmark). Fluorescence was visualized using an Olympus Fluoview FV1000 laser confocal microscope. Images are the product of 3-fold line averaging. Immunofluorescence experiment using preadipocytes were described previously ^18, 52, 53^. The following antibodies were used: anti-GM130 (1:250; Transduction Lab, Lexington, KY, USA, #610823), anti-Calreticulin (1:250; Thermo Fisher Scientific, #PA3-900), anti-HA (1:250; BioLegend, San Diego, CA, USA, #901513) or anti-HA (1:100; MBL, Tokyo, Japan, #561).

### Gene expression analysis

Total RNA was isolated from tissues using QIAzol (Qiagen, Valencia, CA, USA) following the manufacturer’s protocol. Reverse transcription reactions were performed using High Capacity cDNA Synthesis Kit (Applied Biosystems, Foster, CA, USA). The sequences of the primers used in this study are shown in Table S1. Quantitative reverse transcriptase Polymerase Chain Reaction (qRT-PCR) was performed with SYBR green-fluorescent dye using an ABI 7500 Fast Real-Time PCR System. Relative mRNA expression was determined by relative standard curve methods using *18S* (for mouse), or *Gapdh* (for human) as an internal control to normalize samples.

### Plasmid construction and virus production

C-terminally HA-tagged plasmids expressing mouse ZIP13 (mZIP13-HA) were described previously^18^. Phoenix packaging cells were transfected using retroviral vectors by lipofection. After 48 h, the viral supernatant was collected and filtered. Preadipocytes were incubated for 6 h with viral supernatant supplemented with 10 μg/mL polybrene. For the construction of 4F2hc-m13 and 4F2hc-m14, 4F2hc and mZIP13 or mZIP14 were fused by PCR and In-Fusion® HD Cloning Kit (Takara Bio, Shiga, Japan), according to the manufacturer’s instructions. In-Fusion PCR primers were designed to share 15 bases of homology with the PCR fragments and vector ends. 4F2hc, ZIP13 and ZIP14 plasmids, which had already been constructed previously ^18, 55, 56^, were used for PCR templates. The PCR fragments were subcloned into the mammalian expression vector pcDNA 3.1(-) at an EcoRV site.

### *In vivo* and *ex vivo* lipolysis

To determine *in vivo* lipolysis, circulating plasma NEFA levels were measured after overnight fasting (12 h). Mice were then given an intraperitoneal (i.p.) injection of 10mg/kg for isoproterenol. Plasma NEFAs were measured 30 min after injection. To determine *ex vivo* lipolysis, the release of NEFA from adipose tissue was measured, as previously described ^57^. Briefly, mice were sacrificed, and each type of adipose tissue was excised. The fat pads were washed in prewarmed 2% bovine serum albumin (BSA, Free fatty acid free)-DMEM/F12 medium. Adipose tissue pieces (20 mg) were incubated in 200 μL 2% BSA-DMEM/F12 medium with 5μM Triacsin C (Sigma) in the presence or absence of 10 μM isoproterenol. The medium was removed and used to measure NEFA release (NEFA kit, WAKO). For protein quantification, fat explants were first incubated in 1 mL extraction solution (chloroform: methanol, 2:1) and incubated for 60 min at 37 °C under vigorous shaking. Then, the tissue pieces were transferred to 500 μL lysis solution (0.3 N NaOH/0.1% sodium dodecyl sulfate (SDS)) and incubated overnight at 55 °C under vigorous shaking. Protein content was determined using pierce 660 nm protein assay reagent (Thermo Fisher Scientific), and BSA as the standard.

### Cellular lipolysis assay

Preadipocytes were incubated with a differentiation induction cocktail, and differentiated for 8 days. Lipolysis was assessed by measuring NEFA release in the culture medium. The NEFA assay was performed according to the manufacturer’s instructions ^57^. Briefly, aliquots of cell culture medium (4μL) were added to 80 μL reagent A, and incubated at 37 °C for 10 min. This was followed by the addition of 160 μL reagent B, and the incubation for another 37 °C for 10 min, and then absorbance was measured at 550 nm. NEFA releases was normalized to total protein content using pierce 660 nm protein assay reagent. For inhibitor treatment, cells were incubated with 50 μM rolipram (WAKO) or 10 μM cilostamide (Cayman) for 2 h.

### cAMP analysis

The cAMP enzyme immunoassay (Cayman) was performed according to the manufacture’s protocol. Briefly, preadipocytes were differentiated into mature adipocytes, and treated with 10 μM isoproterenol, and then cells were washed twice with ice-cold phosphate-buffer saline (PBS), lysed by the addition of 1 mL 0.1 M HCl, and centrifuged at 1,000 *g* for 10 min. The supernatant was diluted in enzyme-linked immunosorbent assay (ELISA) buffer (sample dilution: x3) and cAMP levels were measured by enzyme immunoassay according to the manufacturer’s protocol. cAMP levels were normalized to protein content, which was determined by assaying the supernatant using the pierce 660 nm protein assay reagent.

### PDE activity assay

PDE activity was measured using the PDE activity assay kit according to the manufacturer’s protocol (Abcam). PDE activity was normalized to protein content, by assaying the cell lysates using pierce 660 nm protein assay reagent.

### Zn^2+^ and Fe^2+^ measurements

Zn^2+^ levels were measured using FluoZin-3 (Invitrogen, #F24194) as described previously ^16^. Cells were plated onto 24-well plates, and washed with Hanks’ Balanced Salt Solution (HBSS) buffer (Sigma-Aldrich) 3 times. The cells were then incubated with 3 μM FluoZin-3 for 1 h, and washed with HBSS buffer 3 times. Fluorescence was recorded at 495 nm excitation and 517 nm emission using EnSpire multimode plate reader (Parkin Elmer). Fe^2+^ levels were measured using FerroOrange (DOJINDO, Kumamoto, Japan, #F374) as described previously ^58^. Briefly, the cells were incubated with 2 μM FerroOrange for 1 h. Fluorescence was recorded at 543 nm excitation and 580 nm emission. For protein determination, cells were lysed with 250 μL lysis solution (0.3N NaOH/0.1% SDS), and incubated at room temperature under vigorous shaking. Protein content was determined using pierce 660 nm protein assay reagent and BSA as a standard.

### cRNA preparation

The oocyte expression vectors were linearized with Hind III and *in-vitro* transcribed to cRNA using the mMESSAGE mMACHINE T7 Transcription Kit (Thermo Fisher Scientific). Synthesized cRNAs were then polyadenylated with the Poly(A) Tailing Kit (Thermo Fisher Scientific) and purified with the MEGAclear Transcription Clean-Up Kit (Thermo Fisher Scientific), according to the manufacture’s protocols.

### Oocyte preparation and transporter expression

The oocyte transport assay was performed as previously reported ^59^. Briefly, adult female *Xenopus laevis* (Watanabe Zoshoku, Japan) were anesthetized with 1.0 to 1.5 mg/mL tricaine (Tokyo Chemical Industry, Japan) and pieces of the ovary were surgically removed. Oocytes were isolated from the ovary and defolliculated with 1.0 to 1.5 mg/mL collagenase A (Roche) in OR2 buffer (82 mM NaCl, 2.5 mM KCl, 1 mM MgCl_2_, 5 mM HEPES; pH 7.6) at room temperature for 1.5 to 2 h. Defolliculated oocytes were washed more than 5 times with OR2 buffer, and then transferred to ND96 buffer (96 mM NaCl, 2 mM KCl, 1.8 mM CaCl_2_, 1 mM MgCl_2_, 5 mM HEPES; pH 7.5). Stage IV to VI oocytes were selected and injected with 18.4 nL (18.4 ng) of purified cRNAs, or equal volumes of RNA-free water using Nanoject II microinjector (Drummond Scientific), and incubated for 2 days at 18 °C in ND96 buffer before their use in the uptake assay.

### Metal uptake assay

Six to eight cRNA-injected oocytes were placed in a 96-well plate, and washed twice with ND96 buffer. Uptake assays were started by replacing the buffer with 120 μL of iron uptake solution (ND 96 buffer containing 100 μM FeCl_2_, 1 - 5 μCi/mL ^55^FeCl_3_, and 1 mM ascorbic acid) or zinc uptake solution (ND96 containing 100 μM ZnSO_4_ and 1 - 2 μCi/mL ^65^ZnCl_2_), and incubated at 22 °C for 240 min. The iron uptake was stopped by replacing the uptake solution with ice-cold ND96 supplemented with 1 mM ascorbic acid. The oocytes were then transferred to a 24-well plate and washed 5 times with the same buffer, solubilized with 0.0625 N NaOH, and radioactivity was measured by liquid scintillation counting. The zinc uptake was stopped by replacing the uptake solution with ice-cold ND96 buffer supplemented with 5 mM EDTA. The oocyte was transferred to 24-well plate and washed 5 times with the same buffer. The amount of zinc in the oocytes was visualized by a Phosphor Imager (Typhoon FLA 9500; GE Healthcare), and quantified by ImageJ software. Each experiment was performed twice on different days.

### Structure prediction

Protein structure prediction of 4F2hc-m13 was performed using locally installed version of ColabFold v1.3.0 ^60^, which is the optimized implementation version of the AlphaFold2. Five models of 4F2hc-m13 were generated using 3 recycles and refined with AMBER force field. The models were visualized and superimposed with the model of mouse ZIP13 (PDB ID: Q8BZH0) deposited in AlphaFold DB using Mol* Viewer (Fig. S8) ^61^. The predicted models were ranked by their pLDDT value (a per-residue confidence score), and the m13 domain of the top-scored model (rank 1) was superimposed with the m13 domains of the other models (Fig. S9).

### ICP-MS

Preadipocyte and mature adipocytes were harvested with methanol and evaporated with centrifugal concentration ^62^. Samples were digested with 0.5 mL of HNO_3_ (Tamapure-AA-100, Tama Chemical Co. Ltd., Kanagawa, Japan) at 180 °C for 20 min in an ETHOS 1 microwave oven (Milestone, Srl, Sorisole, Italy) and then diluted with ultra-pure water (manufactured by PURELAB Option-R 7 and PURELAB flex UV, Veolia Water Solution and Technologies, Paris, France) to a 5-mL volume. Concentrations of Fe and Zn were determined using inductively coupled plasma-sector field mass spectrometry ICP-SFMS (ELEMENT XR, Thermo Fisher Scientific Inc., Bremen, Germany). ^56^Fe and ^66^Zn were measured using medium resolution mode (R = 4,000). The methanol blank samples were prepared without cells and analyzed in the same manner. The amounts of Fe and Zn amounts were corrected by subtracting the mean amounts of the blank samples.

### Ion binding site prediction and analysis

The 3D structure of mouse ZIP13 (ZIP13-WT), obtained from the AlphaFold2 database (UniProt ID: Q8BZH0) ^63^, served as the structure for ion binding prediction. Structural pairwise alignment between ZIP8, sourced from a previous study ^30,^ and ZIP13, alongside the ZincBind web server ^64^, facilitated the prediction and confirmation of two ion coordination sites denoted as M1 and M2. Subsequently, homo Zn^2+^ ions were positioned within M1 and M2 in ZIP13-WT. This methodology was used for the homo Fe^2+^ system preparation. The parameters of the modified 12-6-4 Lennard-Jones (m-12-6-4-LJ) non-bonded model for Zn^2+^ and Fe^2+^ were adopted based on previous recommendations ^30^. The previous m-12-6-4-LJ model was developed to incorporate the polarizabilities of coordinating atoms, which is crucial for accurately capturing the induced dipole effect in MD simulations of ion transport involving metal ions and coordinating atoms ^65-67^. This model was also validated for its effectiveness in simulating metal binding at the selectivity filter, highlighting its sensitivity to the polarizability of coordinating atoms.

Both ZIP13-WT complexes with Zn^2+^ and Fe^2+^ were simulated for 0.5 μs-MD trajectories with 3 replications to observe ion binding stability using AMBER20 package software ^68^, as a standard protocol ^69, 70^. These systems were prepared for 0.5 μs-MD simulations via CHARMM-GUI ^71^, featuring box dimensions of 99 × 99 × 123 Å, with 131 1,2-dilauroyl-sn-glycero-3-phosphocholine (DLPC) molecules per leaflet, and 27,442 TIP3P water molecules. System neutralization was achieved with a 0.15 M KCl concentration, with an additional 0.05 M ZnCl and FeCl added to ZIP13/Zn^2+^ and ZIP13/Fe^2+^, respectively. Protein and ions were treated by ff19SB Amber forcefield ^72^ and 12-6-4 Lennard-Jones model, respectively. Five-step minimization, utilizing steepest descent and conjugate gradient methods with restriction from the protein backbone to the side chain, was conducted for 10,000 steps at each stage. Subsequently, a 36 ns NVT heating phase gradually increased the temperature from 0 to 300 K. Production involved 0.5 μs-MD simulations. These MD trajectories were analyzed to assess ion binding stability by the “hbond” action command implemented in the CPPTRAJ ^73^ module within the AmberTools 21 package program ^74^. This technique resembles traditional hydrogen bond interaction analysis but with the angle cutoff disabled and a distance cutoff set at 3.0 Å. The evaluation of ion binding stability was based on the lifetime and average number of amino acids in contact with the ions.

### Statistical analysis

All quantitative data were reported as the mean ± standard error of the mean (SEM). Statistical differences between the two groups were analyzed by the two-tailed unpaired Student’s *t*-test. For multiple comparisons, analysis of variance (ANOVA) was performed followed by two-way ANOVA with the Bonferroni’s multiple comparison test, or one-way ANOVA with the post-hoc Tukey-Kramer test for comparison among groups, or the post-hoc Dunnett test for comparison of groups with a specific control. A *p*-value of less than 0.05 was considered to indicate a statistically significant difference between groups.

### Ethics approval

All animal experiments were performed in accordance with institutional and national guidelines and regulations and RIKEN Regulations for the Animal Experiments. The study protocol was approved by the Institutional Animal Care and Use Committee of Gunma University (study approval no.: #19-025), and of RIKEN Kobe Branch (Approval number: A2001-03). This study was reported in accordance with ARRIVE guidelines.

## Abbreviations

RER: respiratory exchange ratio
HFD: high-fat-diet
STD: standard diet
ER: endoplasmic reticulum
PKA: protein kinase A
ICP-MS: inductively coupled plasma mass spectrometry
EDSSPD3: Ehlers-Danlos syndrome spondylocheirodysplastic type 3
NEFA: nonesterified fatty acid
MT: metallothionein
PDE: phosphodiesterase
4F2hc: 4F2 cell-surface antigen heavy chain
SVF: stromal vascular fraction
IRP2: iron regulatory protein 2
TfR: transferrin receptor
Fth1: ferritin heavy chain
FTL: ferritin light chain
NAC: N-acetylcysteine
FACS: fluorescence-activated cell sorting
WAT: white adipose tissue.

## Acknowledgments

We thank Dr. H. A. Popiel, Dr. I. Yanatori, Professor K. Ishimori, Professor T. Kitamura, Professor Y. Tamura, Professor S. Toyokuni, Professor J. Shirakawa, Professor S. Nagamori, Professor K. Iwai, and Dr. T. Kambe for their helpful suggestions and discussions, and Ms. Y. Miyazaki, Ms. Y. Tamura, Ms. W. Mizutani, and Ms. A. Suda for their technical assistance. We also appreciate the assistance from the Mouse Facility Core of Gunma University. This work was supported by a Grant-in-Aid for Scientific Research on Innovative Areas “Integrated Bio-metal Science” (MEXT KAKENHI grant number JP22H04801 to A.F.), and JP22K8618, and JP19K08971 (to A.F.), 20H03409 (to T.F.) from the Ministry of Education, Culture, Sports, Science, and Technology, Japan; grants from Takeda Science Foundation, The Koyanagi Foundation, The TANITA Healthy Weight Community Trust, Kanae Foundation for the promotion of Medical Science, Front Runner of Future Diabetes Research Foundation, the Shiseido Female Researcher Science Grant, The Cell Science Research Foundation, and The Ichiro Kanehara Foundation (all to A.F.). This work was also supported by research grant from the Secom Science and Technology Foundation (to Y.F.). This work was carried out by the joint research program of the Institute for Molecular and Cellular Regulation, Gunma University (18010 to T.F.).

## Author contributions

A.F. conceived the project. A.F. performed the experiments and analyzed the data. G.T., and T.K. performed the oocyte transport experiments. A.F., and D.S. conducted the metabolic studies. K.H. performed *in silico* study of ion binding site evaluation. T.I., and I.A. performed the ICP-MS and fluorescence-activated cell sorting (FACS) experiments, respectively. M. Shimura, T.S., and Y.N. analyzed the data. M. Shigeta, H.K., and T.F. generated *Zip13*-flox mice. T.H., and I.H. provided the adiponectin-Cre mice. A.F. wrote the original manuscript, and A.F., H. Shibata, K.H., Y.S., H Sakurai, S.K., H.W., T.F., and Y.F. reviewed the manuscript. T.F., and Y.F. supervised the project.

## References

1 Auger, C. & Kajimura, S. Adipose Tissue Remodeling in Pathophysiology. Annu Rev Pathol 18, 71–93 (2023). 10.1146/annurev-pathol-042220-023633

2 Angueira, A. R. et al. Defining the lineage of thermogenic perivascular adipose tissue. Nat Metab 3, 469–484 (2021). 10.1038/s42255-021-00380-0

3 Tamaki, M. et al. The diabetes-susceptible gene SLC30A8/ZnT8 regulates hepatic insulin clearance. J Clin Invest 123, 4513–4524 (2013). 10.1172/JCI68807

4 Krishnamoorthy, L. et al. Copper regulates cyclic-AMP-dependent lipolysis. Nat Chem Biol 12, 586–592 (2016). 10.1038/nchembio.2098

5 Joffin, N. et al. Adipose tissue macrophages exert systemic metabolic control by manipulating local iron concentrations. Nat Metab 4, 1474–1494 (2022). 10.1038/s42255-022-00664-z

6 Misu, H. et al. Deficiency of the hepatokine selenoprotein P increases responsiveness to exercise in mice through upregulation of reactive oxygen species and AMP-activated protein kinase in muscle. Nat Med 23, 508–516 (2017). 10.1038/nm.4295

7 Jiang, J. et al. Thermogenic adipocyte-derived zinc promotes sympathetic innervation in male mice. Nat Metab 5, 481–494 (2023). 10.1038/s42255-023-00751-9

8 Fukunaka, A. & Fujitani, Y. Role of Zinc Homeostasis in the Pathogenesis of Diabetes and Obesity. Int J Mol Sci 19 (2018). 10.3390/ijms19020476

9 Jeong, J. & Eide, D. J. The SLC39 family of zinc transporters. Mol Aspects Med 34, 612–619 (2013). 10.1016/j.mam.2012.05.011

10 Jenkitkasemwong, S. et al. SLC39A14 Is Required for the Development of Hepatocellular Iron Overload in Murine Models of Hereditary Hemochromatosis. Cell Metab 22, 138–150 (2015). 10.1016/j.cmet.2015.05.002

11 Wang, G. et al. Metastatic cancers promote cachexia through ZIP14 upregulation in skeletal muscle. Nat Med 24, 770–781 (2018). 10.1038/s41591-018-0054-2

12 Lin, W. et al. Hepatic metal ion transporter ZIP8 regulates manganese homeostasis and manganese-dependent enzyme activity. J Clin Invest 127, 2407–2417 (2017). 10.1172/JCI90896

13 Park, J. H. et al. SLC39A8 Deficiency: A Disorder of Manganese Transport and Glycosylation. Am J Hum Genet 97, 894–903 (2015). 10.1016/j.ajhg.2015.11.003

14 Scheiber, I. F., Wu, Y., Morgan, S. E. & Zhao, N. The intestinal metal transporter ZIP14 maintains systemic manganese homeostasis. J Biol Chem 294, 9147–9160 (2019). 10.1074/jbc.RA119.008762

15 Jeong, J. et al. Promotion of vesicular zinc efflux by ZIP13 and its implications for spondylocheiro dysplastic Ehlers-Danlos syndrome. Proc Natl Acad Sci U S A 109, E3530–3538 (2012). 10.1073/pnas.1211775110

16 Bin, B. H. et al. Biochemical characterization of human ZIP13 protein: a homo-dimerized zinc transporter involved in the spondylocheiro dysplastic Ehlers-Danlos syndrome. J Biol Chem 286, 40255–40265 (2011). 10.1074/jbc.M111.256784

17 Bin, B. H. et al. Molecular pathogenesis of spondylocheirodysplastic Ehlers-Danlos syndrome caused by mutant ZIP13 proteins. EMBO Mol Med 6, 1028–1042 (2014). 10.15252/emmm.201303809

18 Fukunaka, A. et al. Zinc transporter ZIP13 suppresses beige adipocyte biogenesis and energy expenditure by regulating C/EBP-beta expression. PLoS Genet 13, e1006950 (2017). 10.1371/journal.pgen.1006950

19 Speliotes, E. K. et al. Association analyses of 249,796 individuals reveal 18 new loci associated with body mass index. Nat Genet 42, 937–948 (2010). 10.1038/ng.686

20 Willer, C. J. et al. Discovery and refinement of loci associated with lipid levels. Nat Genet 45, 1274–1283 (2013). 10.1038/ng.2797

21 Fukada, T. et al. The zinc transporter SLC39A13/ZIP13 is required for connective tissue development; its involvement in BMP/TGF-beta signaling pathways. PLoS One 3, e3642 (2008). 10.1371/journal.pone.0003642

22 Giunta, C. et al. Spondylocheiro dysplastic form of the Ehlers-Danlos syndrome--an autosomal-recessive entity caused by mutations in the zinc transporter gene SLC39A13. Am J Hum Genet 82, 1290–1305 (2008). 10.1016/j.ajhg.2008.05.001

23 Xiao, G., Wan, Z., Fan, Q., Tang, X. & Zhou, B. The metal transporter ZIP13 supplies iron into the secretory pathway in Drosophila melanogaster. Elife 3, e03191 (2014). 10.7554/eLife.03191

24 Hirayama T, et al. A highly selective turn-on fluorescent probe for iron (II) to visualize labile iron in living cells. Chemical Science 3, (2013). 10.1039/C2SC21649C

25 Zhang, T. et al. Crystal structures of a ZIP zinc transporter reveal a binuclear metal center in the transport pathway. Sci Adv 3, e1700344 (2017). 10.1126/sciadv.1700344

26 Rumberger, J. M., Peters, T., Jr., Burrington, C. & Green, A. Transferrin and iron contribute to the lipolytic effect of serum in isolated adipocytes. Diabetes 53, 2535–2541 (2004). 10.2337/diabetes.53.10.2535

27 Grabner, G. F., Xie, H., Schweiger, M. & Zechner, R. Lipolysis: cellular mechanisms for lipid mobilization from fat stores. Nat Metab 3, 1445–1465 (2021). 10.1038/s42255-021-00493-6

28 Snyder, P. B., Esselstyn, J. M., Loughney, K., Wolda, S. L. & Florio, V. A. The role of cyclic nucleotide phosphodiesterases in the regulation of adipocyte lipolysis. J Lipid Res 46, 494–503 (2005). 10.1194/jlr.M400362-JLR200

29 Makino, H., de Buschiazzo, P. M., Pointer, R. H., Jordan, J. E. & Kono, T. Characterization of insulin-sensitive phosphodiesterase in fat cells. I. Effects of salts and oxidation-reduction agents. J Biol Chem 255, 7845–7849 (1980).

30 Jiang, Y. et al. Rational engineering of an elevator-type metal transporter ZIP8 reveals a conditional selectivity filter critically involved in determining substrate specificity. Commun Biol 6, 778 (2023). 10.1038/s42003-023-05146-w

31 Harrison, A. V., Lorenzo, F. R. & McClain, D. A. Iron and the Pathophysiology of Diabetes. Annu Rev Physiol 85, 339–362 (2023). 10.1146/annurev-physiol-022522-102832

32 Iwasaki, T. et al. Serum ferritin is associated with visceral fat area and subcutaneous fat area. Diabetes Care 28, 2486–2491 (2005). 10.2337/diacare.28.10.2486

33 Cepeda-Lopez, A. C. et al. Sharply higher rates of iron deficiency in obese Mexican women and children are predicted by obesity-related inflammation rather than by differences in dietary iron intake. Am J Clin Nutr 93, 975–983 (2011). 10.3945/ajcn.110.005439

34 Pinhas-Hamiel, O. et al. Greater prevalence of iron deficiency in overweight and obese children and adolescents. Int J Obes Relat Metab Disord 27, 416–418 (2003). 10.1038/sj.ijo.0802224

35 Cohen, P. & Kajimura, S. The cellular and functional complexity of thermogenic fat. Nat Rev Mol Cell Biol 22, 393–409 (2021). 10.1038/s41580-021-00350-0

36 Sakers, A., De Siqueira, M. K., Seale, P. & Villanueva, C. J. Adipose-tissue plasticity in health and disease. Cell 185, 419–446 (2022). 10.1016/j.cell.2021.12.016

37 Long, J. Z. et al. A smooth muscle-like origin for beige adipocytes. Cell Metab 19, 810–820 (2014). 10.1016/j.cmet.2014.03.025

38 Oguri, Y. et al. CD81 Controls Beige Fat Progenitor Cell Growth and Energy Balance via FAK Signaling. Cell 182, 563–577 e520 (2020). 10.1016/j.cell.2020.06.021

39 Abe, I. et al. Lipolysis-derived linoleic acid drives beige fat progenitor cell proliferation. Dev Cell 57, 2623–2637 e2628 (2022). 10.1016/j.devcel.2022.11.007

40 Xu, J., Wan, Z. & Zhou, B. Drosophila ZIP13 is posttranslationally regulated by iron-mediated stabilization. Biochim Biophys Acta Mol Cell Res 1866, 1487–1497 (2019). 10.1016/j.bbamcr.2019.06.009

41 Zhao, M. & Zhou, B. A distinctive sequence motif in the fourth transmembrane domain confers ZIP13 iron function in Drosophila melanogaster. Biochim Biophys Acta Mol Cell Res 1867, 118607 (2020). 10.1016/j.bbamcr.2019.118607

42 Joffin, N. et al. Mitochondrial metabolism is a key regulator of the fibro-inflammatory and adipogenic stromal subpopulations in white adipose tissue. Cell Stem Cell 28, 702–717 e708 (2021). 10.1016/j.stem.2021.01.002

43 Moreno-Navarrete, J. M., Ortega, F., Moreno, M., Ricart, W. & Fernandez-Real, J. M. Fine-tuned iron availability is essential to achieve optimal adipocyte differentiation and mitochondrial biogenesis. Diabetologia 57, 1957–1967 (2014). 10.1007/s00125-014-3298-5

44 Pang, C., Chai, J., Zhu, P., Shanklin, J. & Liu, Q. Structural mechanism of intracellular autoregulation of zinc uptake in ZIP transporters. Nat Commun 14, 3404 (2023). 10.1038/s41467-023-39010-6

45 Zhang, Y. et al. Structural insights into the elevator-type transport mechanism of a bacterial ZIP metal transporter. Nat Commun 14, 385 (2023). 10.1038/s41467-023-36048-4

46 Wiuf, A. et al. The two-domain elevator-type mechanism of zinc-transporting ZIP proteins. Sci Adv 8, eabn4331 (2022). 10.1126/sciadv.abn4331

47 Gaither, L. A. & Eide, D. J. Functional expression of the human hZIP2 zinc transporter. J Biol Chem 275, 5560–5564 (2000). 10.1074/jbc.275.8.5560

48 Hoch, E., Levy, M., Hershfinkel, M. & Sekler, I. Elucidating the H(+) Coupled Zn(2+) Transport Mechanism of ZIP4; Implications in Acrodermatitis Enteropathica. Int J Mol Sci 21 (2020). 10.3390/ijms21030734

49 Shigihara, N. et al. Human IAPP-induced pancreatic beta cell toxicity and its regulation by autophagy. J Clin Invest 124, 3634–3644 (2014). 10.1172/JCI69866

50 Yagi, T. et al. A novel ES cell line, TT2, with high germline-differentiating potency. Anal Biochem 214, 70-76 (1993). 10.1006/abio.1993.1458

51 Hausman, D. B., Park, H. J. & Hausman, G. J. Isolation and culture of preadipocytes from rodent white adipose tissue. Methods Mol Biol 456, 201–219 (2008). 10.1007/978-1-59745-245-8_15

52 Fukunaka, A. et al. Tissue nonspecific alkaline phosphatase is activated via a two-step mechanism by zinc transport complexes in the early secretory pathway. J Biol Chem 286, 16363–16373 (2011). 10.1074/jbc.M111.227173

53 Fukunaka, A. et al. Demonstration and characterization of the heterodimerization of ZnT5 and ZnT6 in the early secretory pathway. J Biol Chem 284, 30798–30806 (2009). 10.1074/jbc.M109.026435

54 Fukunaka, A. et al. Zinc and iron dynamics in human islet amyloid polypeptide-induced diabetes mouse model. Sci Rep 13, 3484 (2023). 10.1038/s41598-023-30498-y

55 Kanai, Y. et al. Expression cloning and characterization of a transporter for large neutral amino acids activated by the heavy chain of 4F2 antigen (CD98). J Biol Chem 273, 23629–23632 (1998). 10.1074/jbc.273.37.23629

56 Hojyo, S. et al. The zinc transporter SLC39A14/ZIP14 controls G-protein coupled receptor-mediated signaling required for systemic growth. PLoS One 6, e18059 (2011). 10.1371/journal.pone.0018059

57 Schweiger, M. et al. Measurement of lipolysis. Methods Enzymol 538, 171–193 (2014). 10.1016/B978-0-12-800280-3.00010-4

58 Hirayama, T. Fluorescent probes for the detection of catalytic Fe(II) ion. Free Radic Biol Med 133, 38–45 (2019). 10.1016/j.freeradbiomed.2018.07.004

59 Jutabha, P. et al. Human sodium phosphate transporter 4 (hNPT4/SLC17A3) as a common renal secretory pathway for drugs and urate. J Biol Chem 285, 35123–35132 (2010). 10.1074/jbc.M110.121301

60 Mirdita, M. et al. ColabFold: making protein folding accessible to all. Nat Methods 19, 679–682 (2022). 10.1038/s41592-022-01488-1

61 David Sehnal, et al. Mol* Viewer: modern web app for 3D visualization and analysis of large biomolecular structures. Nucleic Acids Res. 49, W431–W437 (2021). 10.1093/nar/gkab314

62 Shimura, M. et al. Imaging of intracellular fatty acids by scanning X-ray fluorescence microscopy. FASEB J 30, 4149–4158 (2016). 10.1096/fj.201600569R

63 Jumper, J., et al. Highly accurate protein structure prediction with AlphaFold. Nature 596, 583–589 (2021). 10.1038/s41586-021-03819-2

64 Ireland, S. M. & Martin, A. C. R. ZincBind-the database of zinc binding sites. Database (Oxford) 2019 (2019). 10.1093/database/baz006

65 Parise, A. & Magistrato, A. Assessing the mechanism of fast-cycling cancer-associated mutations of Rac1 small Rho GTPase. Protein Sci 33, e4939 (2024). 10.1002/pro.4939

66 Puyo-Fourtine, et al. Consistent Picture of Phosphate–Divalent Cation Binding from Models with Implicit and Explicit Electronic Polarization. The Journal of Physical Chemistry B 126, 4022–4034 (2022). 10.1021/acs.jpcb.2c01158

67 Mels, O. et al. Benchmarking biomolecular force field-based Zn2+ for mono- and bimetallic ligand binding sites. Journal of Computational Chemistry 44, 912–926 (2023). 10.1002/jcc.27052

68 D.A. Case, K. B., I.Y. Ben-Shalom, S.R. Brozell, D.S. Cerutti, T.E. Cheatham, III, V.W.D. Cruzeiro, T.A. Darden, R.E. Duke, G. Giambasu, M.K. Gilson, H. Gohlke, A.W. Goetz, R. Harris, S. Izadi, S.A. Izmailov, K. Kasavajhala, A. Kovalenko, R. Krasny, T. Kurtzman, T.S. Lee, S. LeGrand, P. Li, C. Lin, J. Liu, T. Luchko, R. Luo, V. Man, K.M. Merz, Y. Miao, O. Mikhailovskii, G. Monard, H. Nguyen, A. Onufriev, F. Pan, S. Pantano, R. Qi, D.R. Roe, A. Roitberg, C. Sagui, S. Schott-Verdugo, J. Shen, C.L. Simmerling, N.R. Skrynnikov, J. Smith, J. Swails, R.C. Walker, J. Wang, L. Wilson, R.M. Wolf, X. Wu, Y. Xiong, Y. Xue, D.M. York and P.A. Kollman. AMBER 2020. (2020).

69 Hengphasatporn, K. et al. Promising SARS-CoV-2 main protease inhibitor ligand-binding modes evaluated using LB-PaCS-MD/FMO. Scientific Reports 12, 17984 (2022). 10.1038/s41598-022-22703-1

70 Hengphasatporn, K. et al. Halogenated Baicalein as a Promising Antiviral Agent toward SARS-CoV-2 Main Protease. Journal of Chemical Information and Modeling 62, 1498–1509 (2022). 10.1021/acs.jcim.1c01304

71 Jo, S., Kim, T., Iyer, V. G. & Im, W. CHARMM-GUI: a web-based graphical user interface for CHARMM. J Comput Chem 29, 1859–1865 (2008). 10.1002/jcc.20945

72 Tian, C. et al. ff19SB: Amino-Acid-Specific Protein Backbone Parameters Trained against Quantum Mechanics Energy Surfaces in Solution. J Chem Theory Comput 16, 528–552 (2020). 10.1021/acs.jctc.9b00591

73 Roe, D. R. & Cheatham, T. E., 3rd. PTRAJ and CPPTRAJ: Software for Processing and Analysis of Molecular Dynamics Trajectory Data. J Chem Theory Comput 9, 3084–3095 (2013). 10.1021/ct400341p

74 Case, D., et al. AmberTools21. University of California: San Francisco, CA, USA (2021).

