## supplement for "ZIP13 regulates lipid metabolism by changing intracellular iron and zinc balance"

### Supporting information

### Supplemental figure legends

**Fig. S1. Adipo-*Zip13*KO mice showed resistance to develop insulin resistance.**

(A) Lean mass (left) and fat mass (right) of 32 to 33-week-old Ctrl and Adipo-*Zip13*KO mice (n = 8-9 for both) fed a STD. (B) Micro-computed tomography (CT) evaluation of visceral fat mass and subcutaneous fat mass of 32 to 33-week-old Ctrl and Adipo-*Zip13*KO mice fed a STD (n = 8 to 9). (C) Blood glucose concentrations were measured during intraperitoneal glucose tolerance test (IPGTT) in 31 to 32-week-old Ctrl (n = 6) and Adipo-*Zip13*KO (n = 9) mice fed a STD. (D) Insulin concentrations related to of the mice analyzed in (C). (E) Insulin tolerance testing of 19-week-old Ctrl (n = 8) and Adipo-*Zip13*KO (n = 14) mice fed a HFD for 13 weeks. In (A, B, C, D, and E), *$p < 0.05$, **$p < 0.01$, by the two-tailed unpaired Student's *t*-test.

**Fig. S2. The phenotype of Adipo-*Zip13*KO mice related to the adipocyte browning.** (A, B) Whole-body oxygen consumption of 20-week-old Ctrl (n = 8) and Adipo-*Zip13*KO (n = 9) mice fed a STD. (A): VO2 trend, (B): VO2 period. (C) Food intake of 21 to 24 week-old Ctrl (n = 6) and Adipo-*Zip13*KO (n = 7) mice fed a STD. (D) Locomotor activity of 20-week-old Ctrl (n = 8) and Adipo-*Zip13*KO (n = 9) mice fed a STD. (E) Rectal temperatures of 21 to 24 week-old Ctrl (n = 6) and Adipo-*Zip13*KO (n = 7) mice fed a STD. (F) Relative mRNA expression of the indicated genes in the subcutaneous fat tissue of 34 week-old Ctrl (n = 6) and Adipo-*Zip13*KO (n = 6) mice fed a STD. (G) Relative mRNA expression of the indicated genes in differentiated cells from subcutaneous fat tissue of Ctrl and Adipo-*Zip13*KO mice treatment with DMSO and Isoproterenol (Iso). (H) Hematoxylin and eosin (H&E) staining of subcutaneous fat and brown fat tissues in Ctrl and Adipo-

*Zip13*KO mice. Scale bars = 100 μm. (I) Whole-body oxygen consumption (VO2) of 24-week-old Ctrl (n = 9) and Adipo-*Zip13*KO (n = 7) mice fed a STD at the indicated temperature. (J) Relative mRNA expression of the indicated genes from the subcutaneous fat tissue of the mice analyzed in (I). Ctrl (n = 9), Adipo-*Zip13*KO mice (n = 7). Data are shown as the mean ± SEM. In (B, C, D, E, F, I, and J), by the two-tailed unpaired Student's *t*-test. In (G), **$p < 0.01$, ***$p < 0.001$, ****$p < 0.0001$, by one-way ANOVA followed by the post-hoc Tukey-Kramer test. ns, not significant.

**Fig. S3. Adipo-*Zip13*KO mice showed reduced RER.**

(A, B) Whole-body RER of 30-week-old Ctrl (n = 8) and Adipo-*Zip13*KO (n = 9) mice fed a STD. (A): RER trend, (B): RER period. (C) Relative mRNA expression of the indicated genes in 34-week-old Ctrl (n = 5) and Adipo-*Zip13*KO (n = 4) mice on a STD. Data are shown as the mean ± SEM. In (B and C), *$p < 0.05$, **$p < 0.01$, by the two-tailed unpaired Student's *t*-test.

**Fig. S4. Hydroxy radical levels of Ctrl and Adipo-*Zip13*KO cells.**

Hydroxy radical levels are upregulated in Adipo-*Zip13*KO cells. Data are shown as the mean ± SEM. **$p < 0.01$ , ****$p < 0.0001$, by one-way ANOVA followed by the post-hoc Dunnett test.

**Fig. S5. PKA signaling is increased in Adipo-*Zip13*KO cells (related Fig. 4).**

Immunoblotting of pPKA, PKA, and GAPDH in Ctrl and Adipo-*Zip13*KO mature adipocyte cells stimulated with DMSO (Basal) or isoproterenol (Iso) (top). Quantification of pPKA and PKA proteins normalized to PKA and GAPDH, respectively (bottom). Data are shown as the mean ± SEM; **$p < 0.01$, by the two-tailed unpaired Student's *t*-test.

**Fig. S6. *Zip13* expression in adipose tissues.**

(A) Relative *Zip13* expression in the SVF and adipocytes. (B) Cell population of $CD81^{+}$ $Ki67^{+}$ cells in adipose progenitor cells. Data are shown as the mean ± SEM. In (A and B), $^{*}p < 0.05$, $^{**}p < 0.01$, by the two-tailed unpaired Student's *t*-test. APC, adipocyte progenitor cells.

**Fig. S7. The strategy of the conditional knockout at the *Zip13* locus.**

The black boxes indicate exons of the *Zip13* gene. Pr DT-A pA is a diphtheria toxin A fragment gene directed by MC1 promoter for negative selection, Pr Puro pA and Pr Neo pA driven by PGK1 promoter are the drug-resistance genes for positive selection in ES cells. The cassettes flanked by F3 or FRT sequences are excised by flippase to generate the Flox allele. KO allele is generated with a tissue-specific Cre mouse. P1, P2, P3, and P4 are primers for genotyping PCR.

**Fig. S8. Superimposition of mZIP13 and 4F2hc-m13 (five models).**

The predicted models of 4F2hc-m13 were superimposed with the transmembrane region of the model of mouse ZIP13 (PDB ID: Q8BZH0) deposited in AlphaFold DB and the RMSD values were calculated using Mol* Viewer.

**Fig. S9. Superimposition of 4F2hc-m13 (rank 1) and other rank models.**

Top ranked predicted model of 4F2hc-m13 was superimposed with the transmembrane regions of m13 domains of the other models. The RMSD values were calculated using Mol* Viewer.

### Supplemental table

### Table S1 Primer sequences

| Gene | Species | Forward primer | Reverse primer |
|---|---|---|---|
| *18S* | Mouse | TTCTGGCCAACGGTCTAGACAAC | CCAGTGGTCTTGGTGTGCTGA |
| *Ucp1* | Mouse | CACCTTCCCGCTGGACACT | CCCTAGGACACCTTTATACCTAATGG |
| *Pgc1* | Mouse | AGCCGTGACCACTGACAACGAG | GCTGCATGGTTCTGAGTGCTAAG |
| *Cidea* | Mouse | ATCACAACTGGCCTGGTTACG | TACTACCCGGTGTCCATTTCT |
| *Cox8* | Mouse | GAACCATGAAGCCAACGACT | GCGAAGTTCACAGTGGTTCC |
| *aP2* | Mouse | ACACCGAGATTTCCTTCAAACTG | CCATCTAGGGTTATGATGCTCTTCA |
| *Zip13* | Mouse | AGGCCCCCAGCAAAGACCCCA | CTTTTTGCTCACAAGGAAGCT |
| *Cpt1b* | Mouse | GTCGCTTCTTCAAGGTCTGG | AAGAAAGCAGCACGTTCGAT |
| *Cpt2* | Mouse | CAGCACAGCATCGTACCCA | TCCCAATGCCGTTCTCAAAAT |
| *Fasn* | Mouse | GAGGTGGTGATAGCCGGTAT | TGGGTAATCCATAGAGCCCAG |
| *CD36* | Mouse | TGCATTTGCCAATGTCTAGC | CCCTCCAGAATCCAGACAAC |
| *ACC* | Mouse | AATGAACGTGCAATCCCATTTG | ACTCCACATTTGCGTAATTGTTG |
| *Scd1* | Mouse | AGGCCTGTACGGGATCATACT | AGAGCGCTGGTCATGTAGTAG |
| *ATGL* | Mouse | TTCGCAATCTCTACCGCCTC | TGGTTCAGTAGGCCATTCCTC |
| *Elovl3* | Mouse | TCCGCGTTCTCATGTAGGTCT | GGACCTGATGCAACCCTATGA |
| *HSL* | Mouse | GCGCTGGAGGAGTGTTTTT | CCGCTCTCCAGTTGAACC |
| *MGL* | Mouse | AGGCGAACTCCACAGAATGTT | ACAAAAGAGGTACTGTCCGTCT |
| *Plin1* | Mouse | CTGTGTGCAATGCCTATGAGA | CTGGAGGGTATTGAAGAGCCG |
| *MT1* | Human | AGAGTGCAAATGCACCTCCTGC | CGGACATCAGGCACAGCAGCT |
| *GAPDH* | Human | CGAGATCCCTCCAAAATCAA | CATGAGTCCTTCCACGATACCAA |

### Supplemental methods

#### Measurement of food intake of mice

Food intake of mice was measured very day for 5 days. The average of food intake of 1 day was shown. During the measurement period, mice were housed in individual cages.

#### Measurement of rectal temperature of mice

Rectal temperature of mice was measured using the D717 pocket-sized thermistor (Takara Thermistor, Yokohama, Japan).

#### Measurement of blood glucose levels and insulin levels of mice

IPGTT and insulin tolerance test (ITT) were performed as described previously [3, 49] . For IPGTT, animals were fasted for 13 h, and then injected i.p. with 2 g/kg glucose. Glucose levels were measured using a glucose analyzer (Glutest Mint, Sanwa Chemical Co., Japan). Insulin levels were measured using an ELISA kit (Morinaga Co., Kanagawa, Japan). For the ITT, mice were injected i.p. with insulin (0.75 units/kg).

#### Body composition analysis using micro-CT

CT was performed under isoflurane anesthesia using a LaTheta micro-CT scanner (Hitachi, Tokyo, Japan). The abdominal region between the first and sixth lumbar vertebrae was scanned, and the sizes of the fat and soft tissue compartments were measured.

**Tissue histology**

For H&E staining, the tissues of mice were fixed in 4% paraformaldehyde overnight at 4 °C, followed by dehydration in 70% ethanol. After the dehydration procedure, tissues were embedded in paraffin, sectioned at a thickness of 5 □m, and stained with H&E following a standard protocol.

**Measurement of hydroxy-radical levels**

Hydroxy-radical levels were measured using hydroxyphenyl fluorescein (HPF) (Goryo Chemical, #FSK3001-01). Cells were plated onto 24-well plates, and washed with Hanks' Balanced Salt Solution (HBSS) buffer (Sigma-Aldrich) 3 times. The cells were then incubated with 10 μM HPF for 30 min, add DMSO or isoproterenol for 30 min, and washed with HBSS buffer 3 times. Fluorescence was recorded at 490 nm excitation and 515 nm emission using EnSpire multimode plate reader (Parkin Elmer). For protein determination, cells were lysed with 250 μL lysis solution (0.3N NaOH/0.1% SDS), and incubated at room temperature under vigorous shaking. Protein content was determined using pierce 660 nm protein assay reagent and BSA as a standard.

**FACS analysis**

SVF were isolated from the inguinal adipose tissue depots of mice using collagenase D (1.5U

/mL) and Dispase II (2.5U/mL) following the procedure. MACS Non-Adipocyte Progenitor Depletion Cocktail for mice (Miltenyi Biotec, USA) and MACS LS columns (Miltenyi Biotec) were used to deplete lineage$^+$ (Lin$^+$) cells. The following procedure was performed for the isolation of mouse CD81$^+$ cells (Lin$^-$: Sca1$^+$:CD81$^+$): Sca-1-PB (1:800, Biolegend, #108120) and CD81-APC (1:50, Biolegend, #104910) in autoMACS Rising Solution (Miltenyi Biotec) containing 0.5% BSA in the dark at 4 °C for 15 min, and incubated in autoMACS Rising Solution containing 0.5% BSA and 0.1% saponin for 15 min and stained with a Ki67-PE antibody (1:300, Biolegend, #151209) for 30 min. Cell population (%) was calculated as the frequency of parent. All the cells were isolated and analyzed using a FACS Aria II equipped with a 100- mm diameter nozzle and CytoFLEX. FlowJo software (version 10.8.1) and CytExpert (version 2.4.0.28) were used for data analysis.

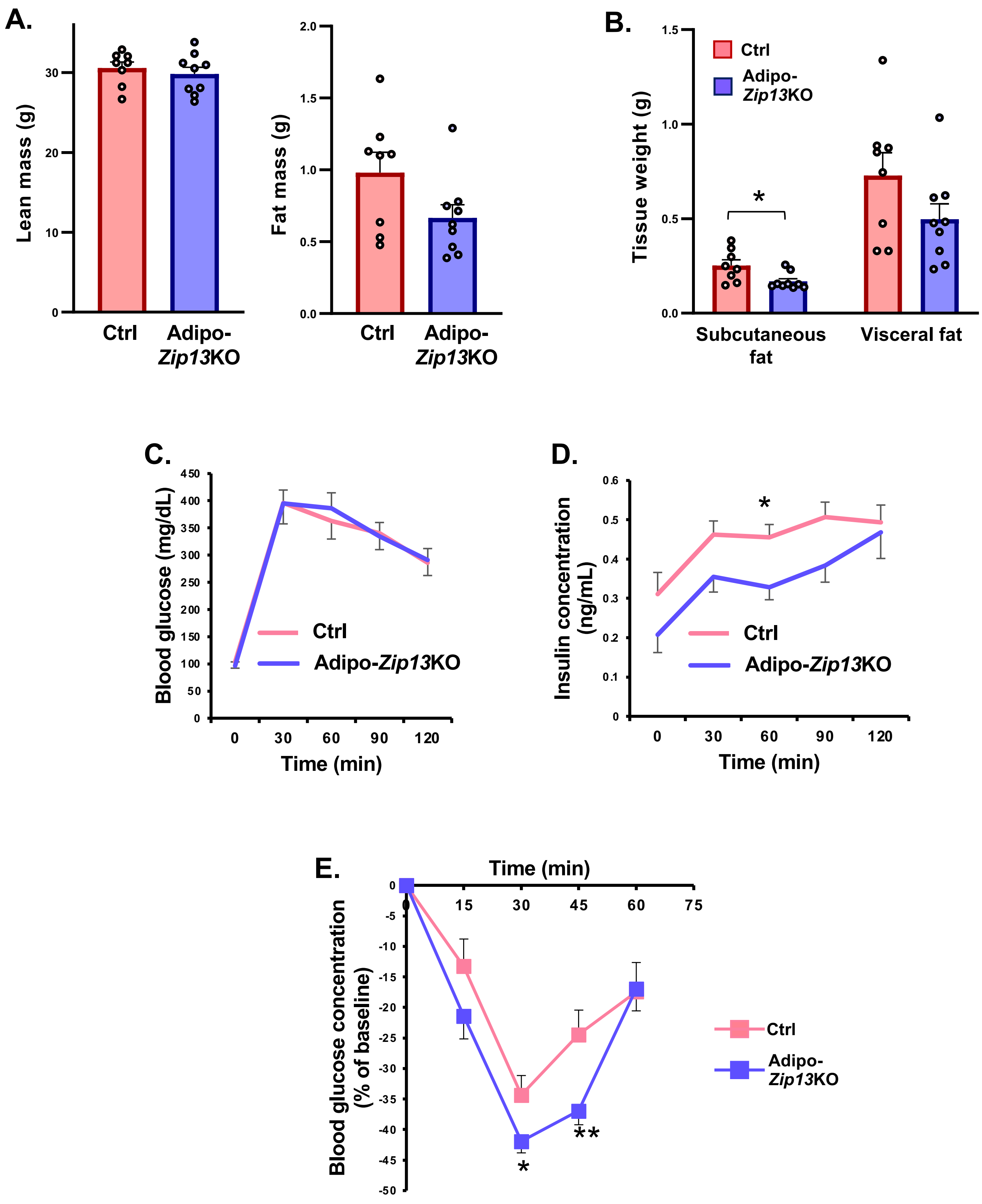


**Fig. S1. Adipo-*Zip13*KO mice showed resistance to develop insulin resistance.**

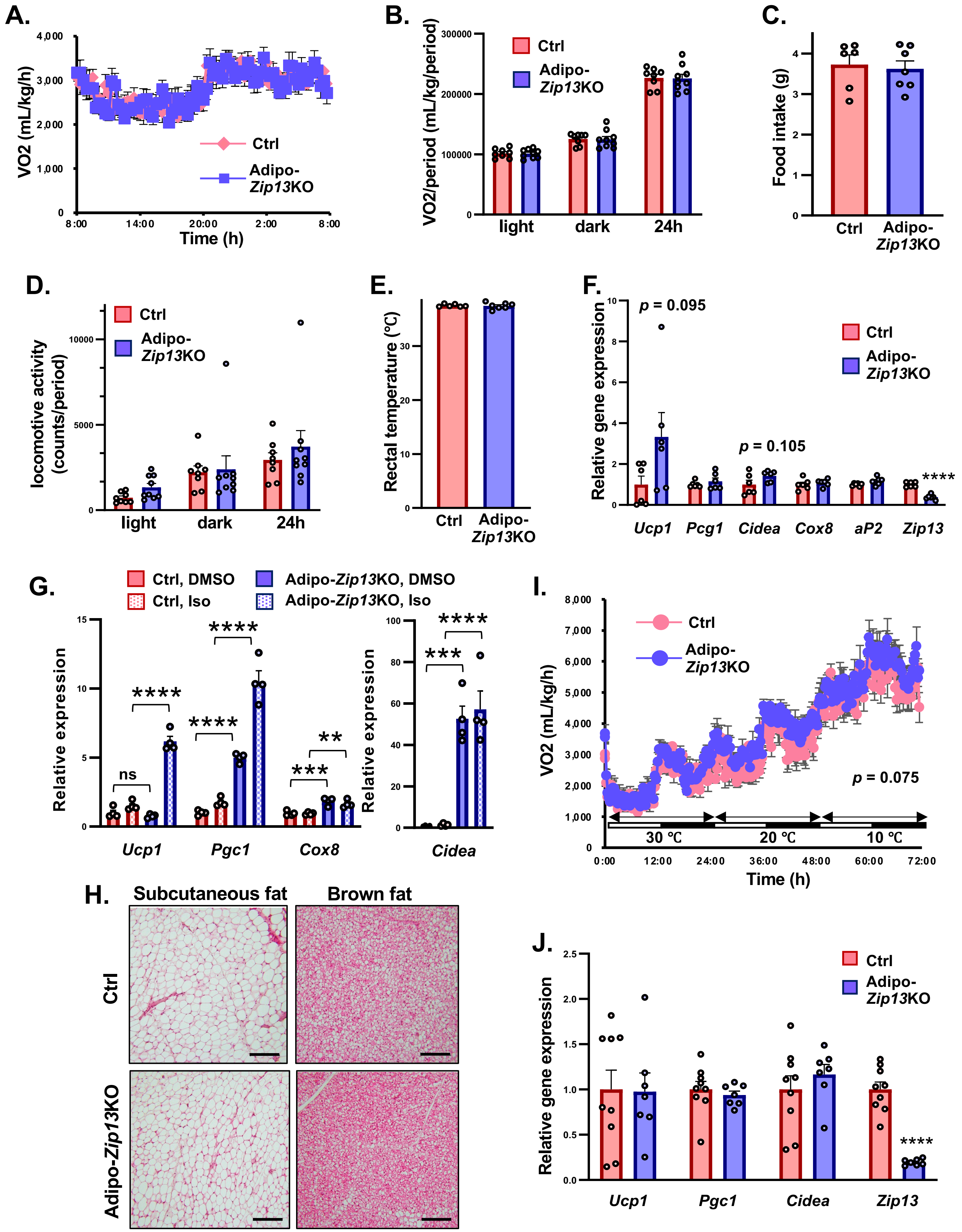


**Fig. S2. The phenotype of Adipo-*Zip13*KO mice related to the adipocyte browning.**

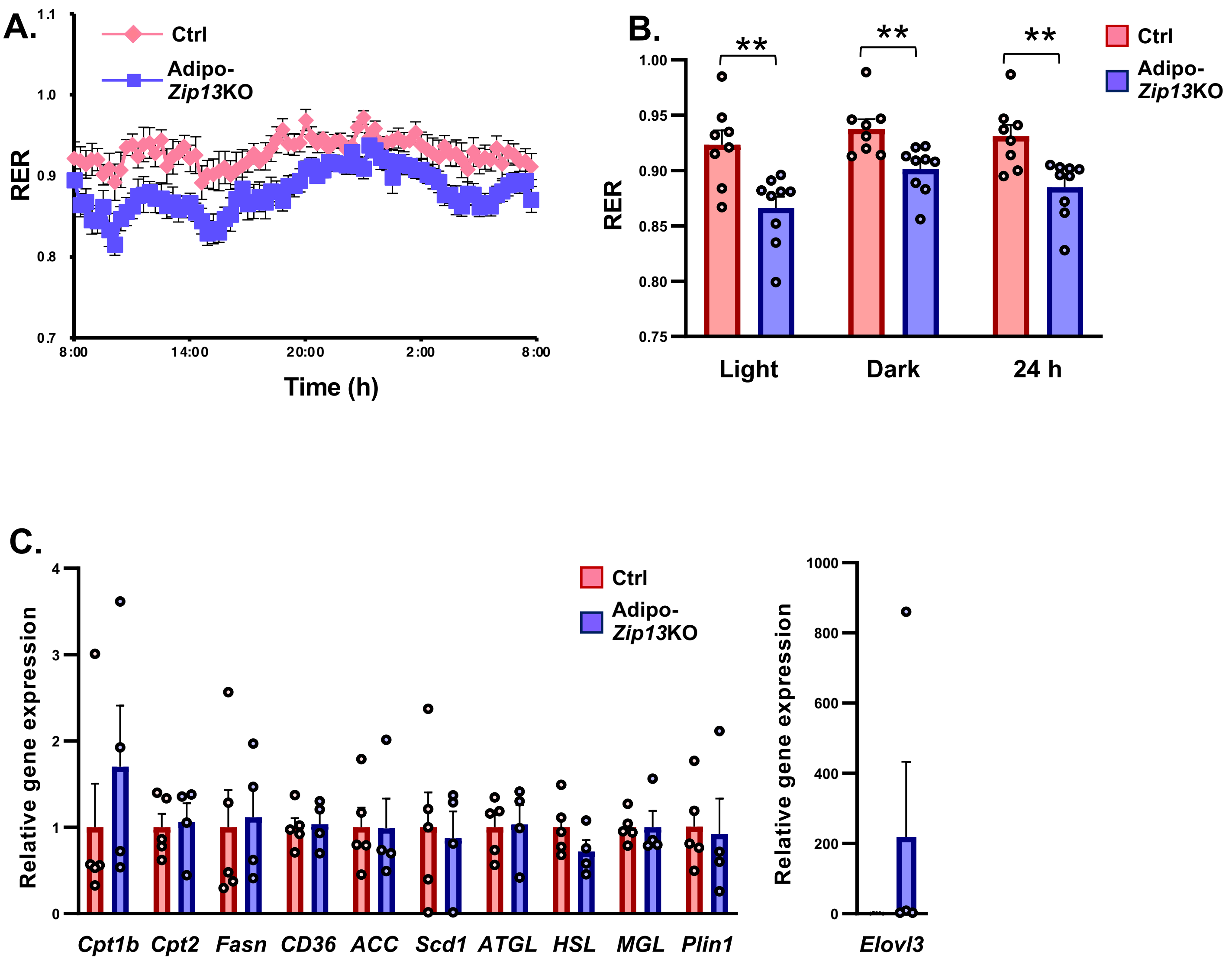


**Fig. S3. Adipo-*Zip13*KO mice showed reduced RER.**

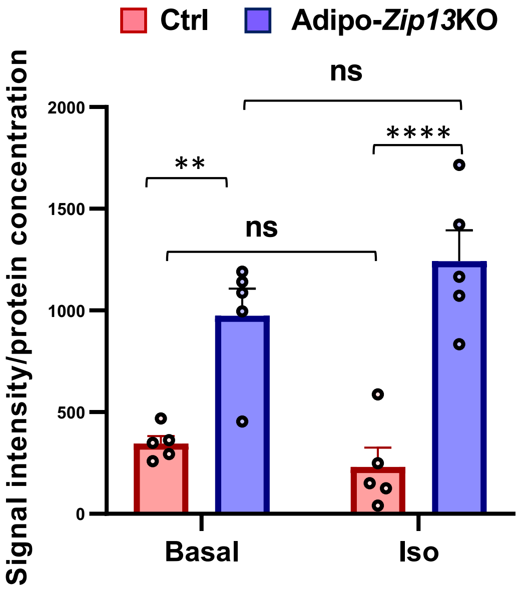


**Fig. S4. Hydroxy radical levels of Ctrl and Adipo-*Zip13*KO cells.**

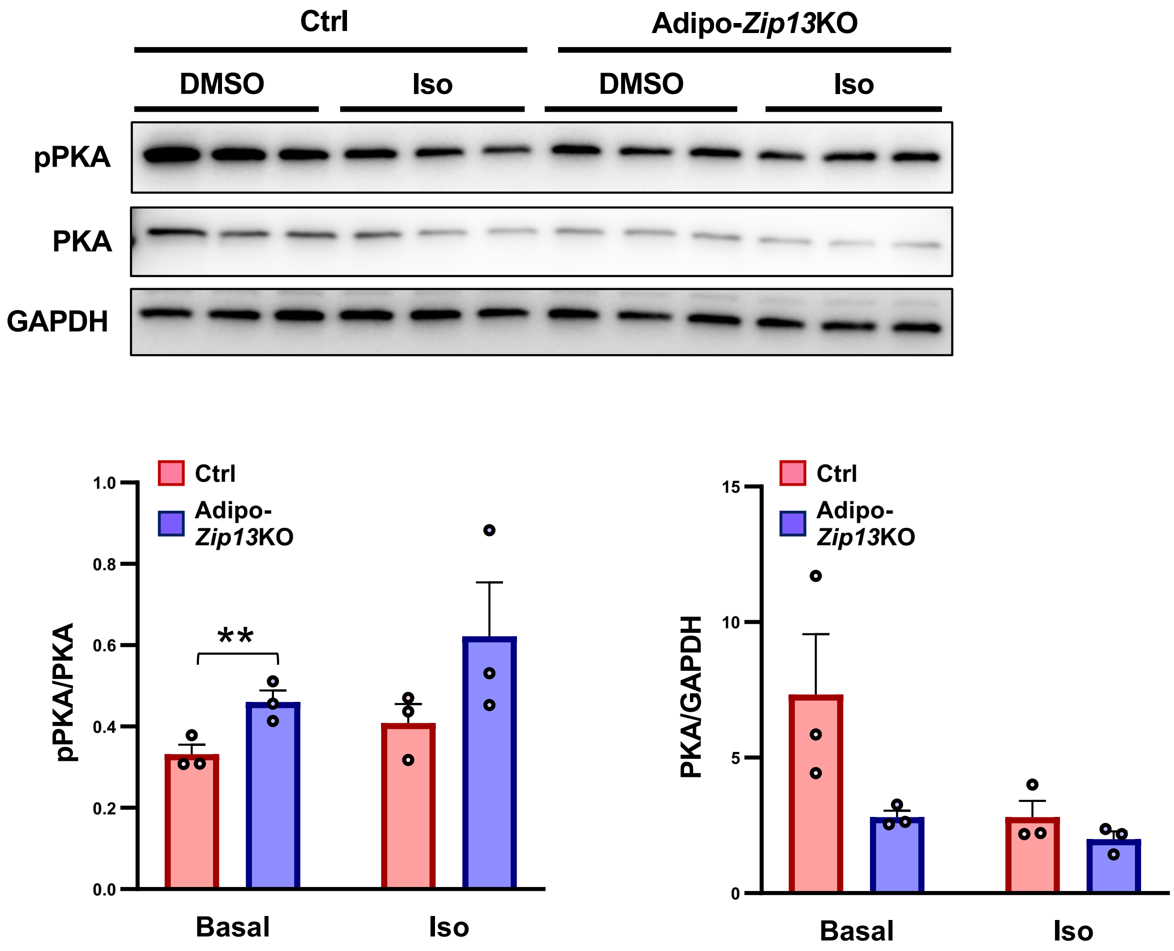


**Fig. S5. PKA signaling is increased in Adipo-*Zip13*KO cells (related Fig. 4).**

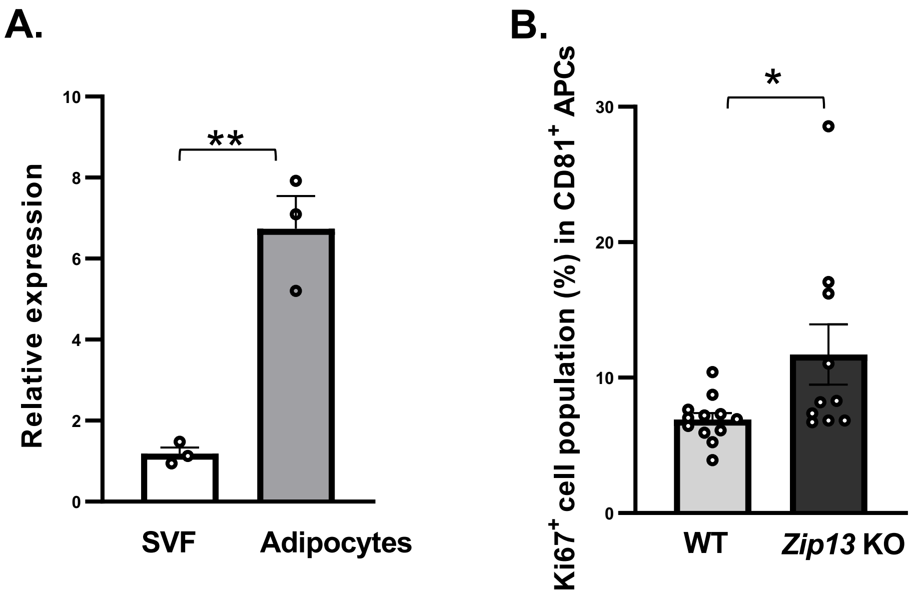


**Fig. S6.** ***Zip13*** **expression in adipose tissues.**

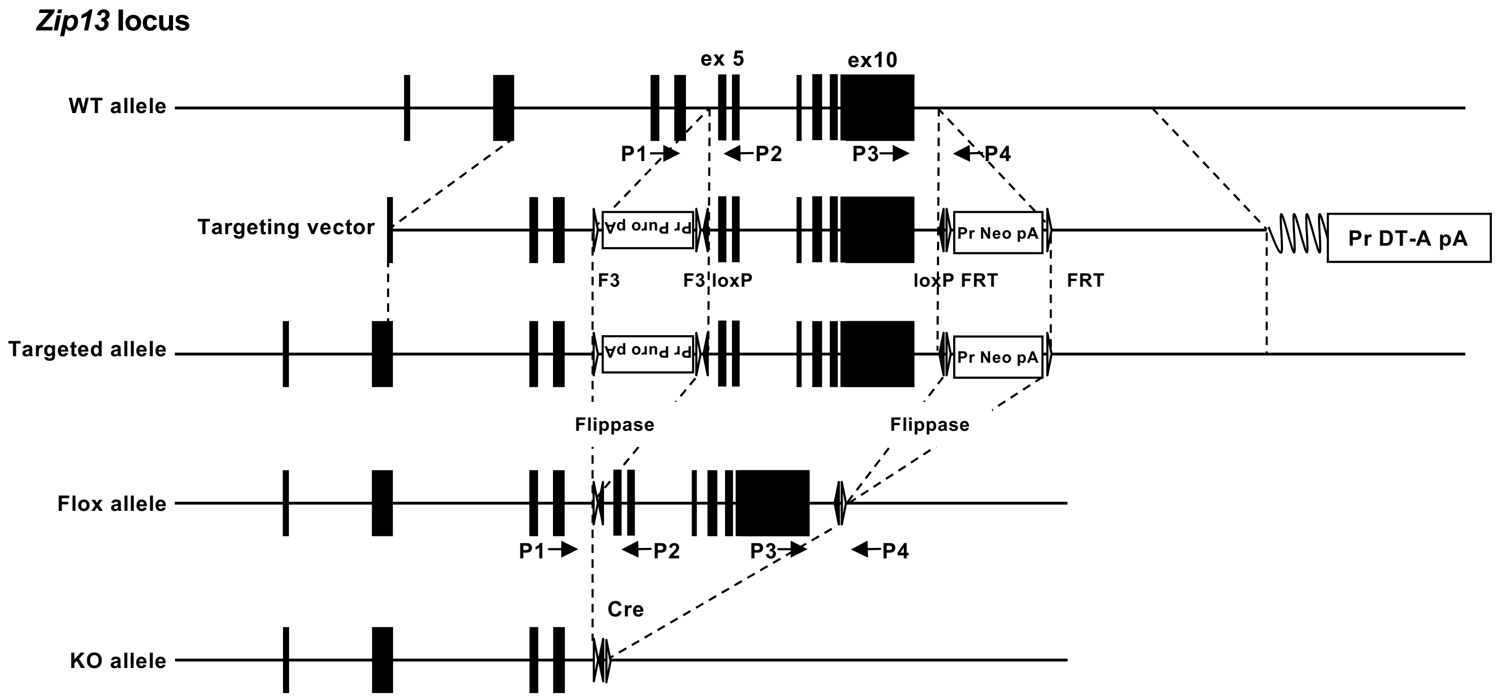


**Fig. S7. The strategy of the conditional knockout at the Z*ip13* locus.**

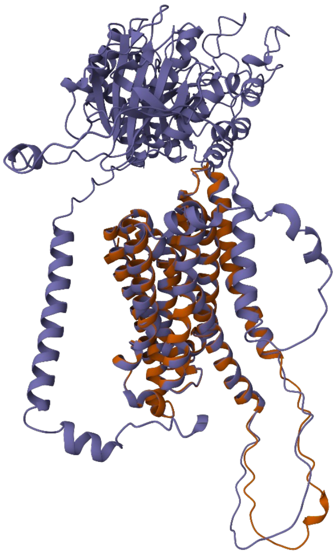

4F2hc-m13 (rank 1)
mZIP13(Q8BZH0)
RMSD: 0.37Å

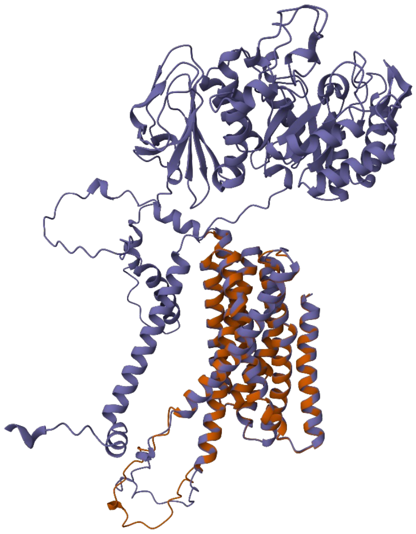

4F2hc-m13 (rank 2)
mZIP13(Q8BZH0)
RMSD: 0.47Å

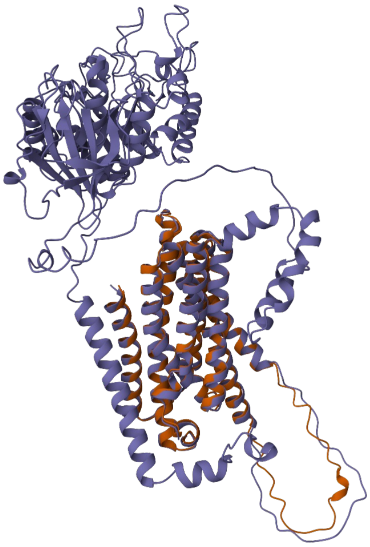

4F2hc-m13 (rank 3)
mZIP13(Q8BZH0)
RMSD: 0.47Å

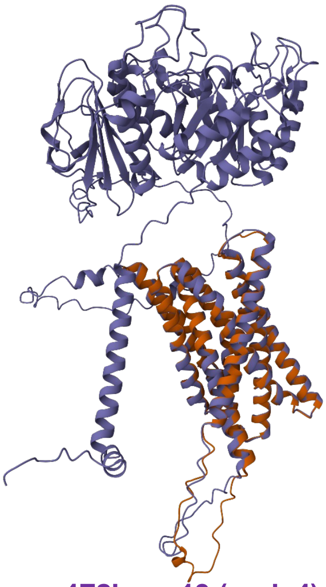

4F2hc-m13 (rank 4)
mZIP13 Q8BZH0)
RMSD: 0.47Å

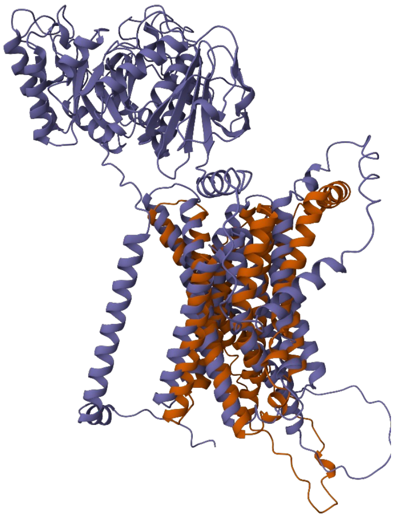

4F2hc-m13 (rank 5)
mZIP13(Q8BZH0)
RMSD: 5.36Å

**Fig. S8. Superimposition of mZIP13 and 4F2hc-m13 (five models).**

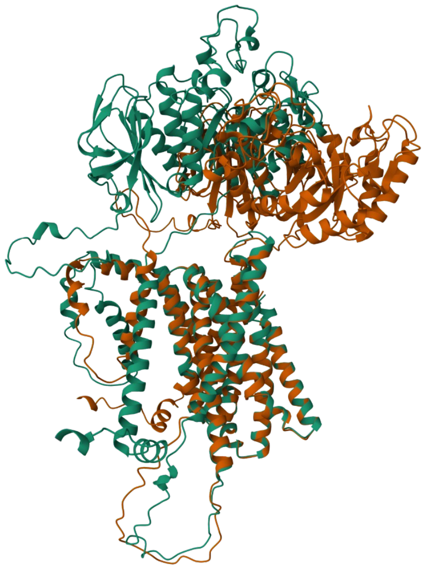

4F2hc-m13 (rank 1)
4F2hc-m13 (rank 2)
RMSD: 0.52Å

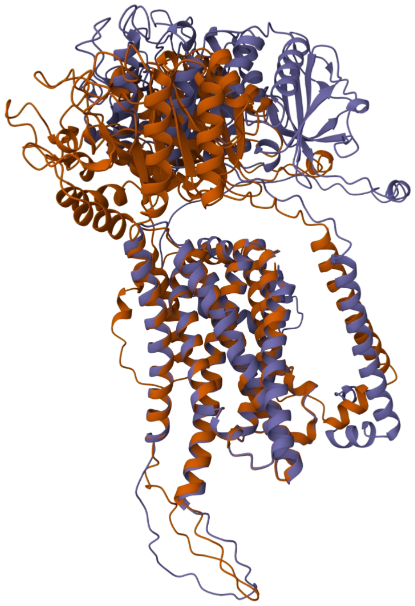

4F2hc-m13 (rank 1)
4F2hc-m13 (rank 3)
RMSD: 0.44 Å

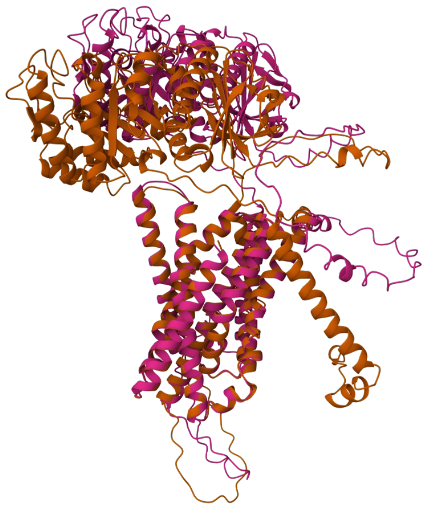

4F2hc-m13 (rank 1)
4F2hc-m13 (rank 4)
RMSD: 0.54Å

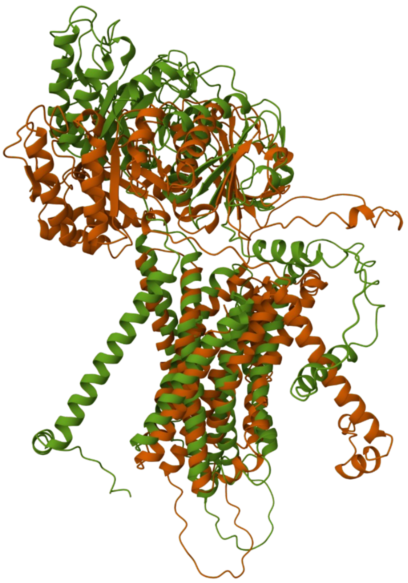

4F2hc-m13 (rank 1)
4F2hc-m13 (rank 5)
RMSD: 5.36Å

**Fig. S9. Superimposition between 4F2hc-m13 (rank1) and other and rank models.**
